# Strain-Level Diversity Decouples Biofilm Architecture, Acidogenic and Aciduric Traits, and Antimicrobial Tolerance in *Streptococcus mutans*

**DOI:** 10.64898/2026.08.25.747077

**Authors:** Kyulim Lee, Daniel I. Peters, Madisen Bangs, Delaney Hancock, Nicole A. Fleming, Jarett Pittman, Taylor S. Martinez, Alyssa N. Deever, Justin R. Kaspar

## Abstract

*Streptococcus mutans* is a key contributor to dental caries, with its capacity to form structured biofilm microcolonies being a principal component of its cariogenic potential. Yet, most mechanistic studies rely on a limited number of laboratory strains and may not capture the functional diversity present across the species. Here, we characterized a panel of phenotypically and genomically diverse *S. mutans* isolates to determine how strain background influences biofilm architecture, extracellular matrix accumulation, acid-associated physiology, environmental responsiveness, and antimicrobial susceptibility. Quantitative high-resolution imaging revealed extensive heterogeneity in produced biofilm microcolony size, structure, and matrix composition, demonstrating that biofilm architecture is not a uniform species-level trait. Interestingly, the commonly used reference strain UA159 displayed an intermediate phenotype related to microcolony size and biofilm organization. Human saliva further altered biofilm structure and matrix accumulation in a strain-dependent manner rather than producing a standard species-wide response. Isolates also differed in growth and retained biofilm biomass under acidic conditions, while acid accumulation within mature biofilms varied independently of average microcolony volume, demonstrating that strains that produce larger microcolonies on average were not necessarily associated with greater acid accumulation. Susceptibility to the antiseptics chlorhexidine and cetylpyridinium chloride likewise differed among isolates and could not be predicted from formed biofilm architecture alone. Together, these findings demonstrate that disease-relevant traits commonly attributed to *S. mutans* are distributed unevenly and only partially coupled across strain backgrounds, with biofilm spatial organization failing to serve as a dominant phenotype linking acid accumulation, acid tolerance, and antimicrobial susceptibility.

## INTRODUCTION

The development of dental caries, a biofilm-mediated disease, is a multifactorial process involving repeated interactions between oral bacteria, dietary sugars and host-related factors such as saliva composition and tooth structure (1). Among the contributors, *Streptococcus mutans* has long been recognized as a key contributor in the initiation of caries due to its efficient metabolism of dietary carbohydrates to organic acids and its pronounced aciduricity, which allows continued growth and acid production even under low pH conditions (2, 3). In addition to these traits, a central determinant of *S. mutans* virulence is its strong capacity to form robust biofilms on the tooth surface, a feature that is critical to its cariogenic potential (2, 4).

Continued advances in imaging technology have reshaped our understanding of how *S. mutans* contribute to cariogenic biofilms *in vivo*. Rather than being uniformly distributed across the tooth surface, *S. mutans* organizes into discrete, rotund microcolony structures embedded within an extracellular polymeric substance (EPS) matrix (5, 6). These microcolonies are spatially organized and enmeshed in EPS, forming diffusion-limited matrices that play a central role in disease pathogenesis by creating acidic microenvironments adjacent to the tooth’s enamel surface (6–9). Within these microcolonies, acids generated through carbohydrate fermentation are retained and concentrated, resulting in sustained low pH conditions that favor enamel demineralization even in the presence of salivary buffering.

The formation and persistence of these microcolonies are driven largely by *S. mutans-* derived glucans, polysaccharides that are synthesized by glucosyltransferases (10), along with extracellular DNA (eDNA) (11). The EPS matrix provides structure cohesion and spatial organization, while imposing diffusion limitations that restrict the penetration of neutralizing agents, including saliva and antimicrobial compounds (5, 12, 13). *In situ* pH mapping and acid stress response analyses have demonstrated that the interiors of *S. mutans* microcolonies remain acidic despite neutral external conditions and are characterized by upregulation of *atpB*, a critical gene for F_1_F_0_-ATPase-mediated proton extrusion and cytoplasmic pH homeostasis (8, 14). Importantly, enzymatic disruption of EPS matrix leads to neutralization of the interiors of the microcolony and marked decrease in *atpB* gene expression, underscoring the matrix’s role in acid pooling and the functional importance of microcolony architecture in cariogenicity (8).

Although previous studies have established the pathogenic relevance of *S. mutans* microcolony formation, the extent to which these phenotypes vary among genetically distinct clinical isolates remains poorly understood. Most mechanistic studies have relied on a limited number of laboratory strains, particularly UA159 (15). However, *S. mutans* possesses an open and variable pangenome, and clinical isolates differ substantially in accessory gene content and in phenotypes such as acid tolerance, oxidative-stress resistance, autolysis, genetic competence, and biofilm formation (16, 17). An earlier phenotypic characterization of a diverse clinical isolate collection revealed considerable variation in these traits, including bulk biofilm-forming capacity (16). However, biofilm formation was assessed primarily by crystal violet (CV) staining, which measures total retained biomass but provides limited insight into three-dimensional architecture, microcolony organization, extracellular matrix composition and spatial distribution, or functional heterogeneity within the biofilm.

To address this gap, we combined quantitative three-dimensional imaging with complementary physiological assays to characterize 20 genomically diverse *S. mutans* isolates. Super-resolution confocal microscopy enabled detailed analysis of strain-specific biofilm architecture, microcolony morphology, spatial organization, and extracellular matrix accumulation. We also examined how each isolate responded to environmental signals, acidic growth conditions, and antimicrobial agents commonly used in oral healthcare. Our findings reveal extensive phenotypic heterogeneity and demonstrate that these disease-relevant traits are only partially coupled across isolates. Thus, behaviors commonly attributed to *S. mutans* at the species level are distributed unevenly among strain backgrounds rather than forming a single coordinated virulence program.

## RESULTS

### Genomically Diverse S. mutans Isolates Form Distinct Microcolony Architectures

To better understand the biofilm formation capacity of *Streptococcus mutans* isolates, we randomly selected a panel of twenty isolates previously characterized for their phenotypic traits (16). Phylogenetic and pangenome analyses confirmed that the panel contains a shared core genome along with substantial differences in accessory gene content **(Figure S1)**.

Biofilms of each isolate were grown for 18 hours in a media (TYGS) consisting of tryptone, yeast extract, 10 mM glucose and 5 mM sucrose to support *S. mutans* glucan production. Biofilms were then imaged using super-resolution confocal microscopy to evaluate biofilm spatial organization, total cell biomass, and individual matrix component accumulation. Microcolony architecture varied significantly across the isolates **(Figure 1A)**. Some isolates formed dense fields of compact, rounded microcolonies, whereas others produced smaller aggregates, diffuse surface-associated biomass, or atypical elongated structures. Quantification of individual microcolony volumes revealed a broad distribution **(Figure 1B)**. Individual microcolony volumes ranged from <100 µm^3^ to >1,000,000 µm^3^, with notable inter-strain variability indicating that each isolate produced a distribution of structures rather than a single characteristic size.

**Figure 1.**
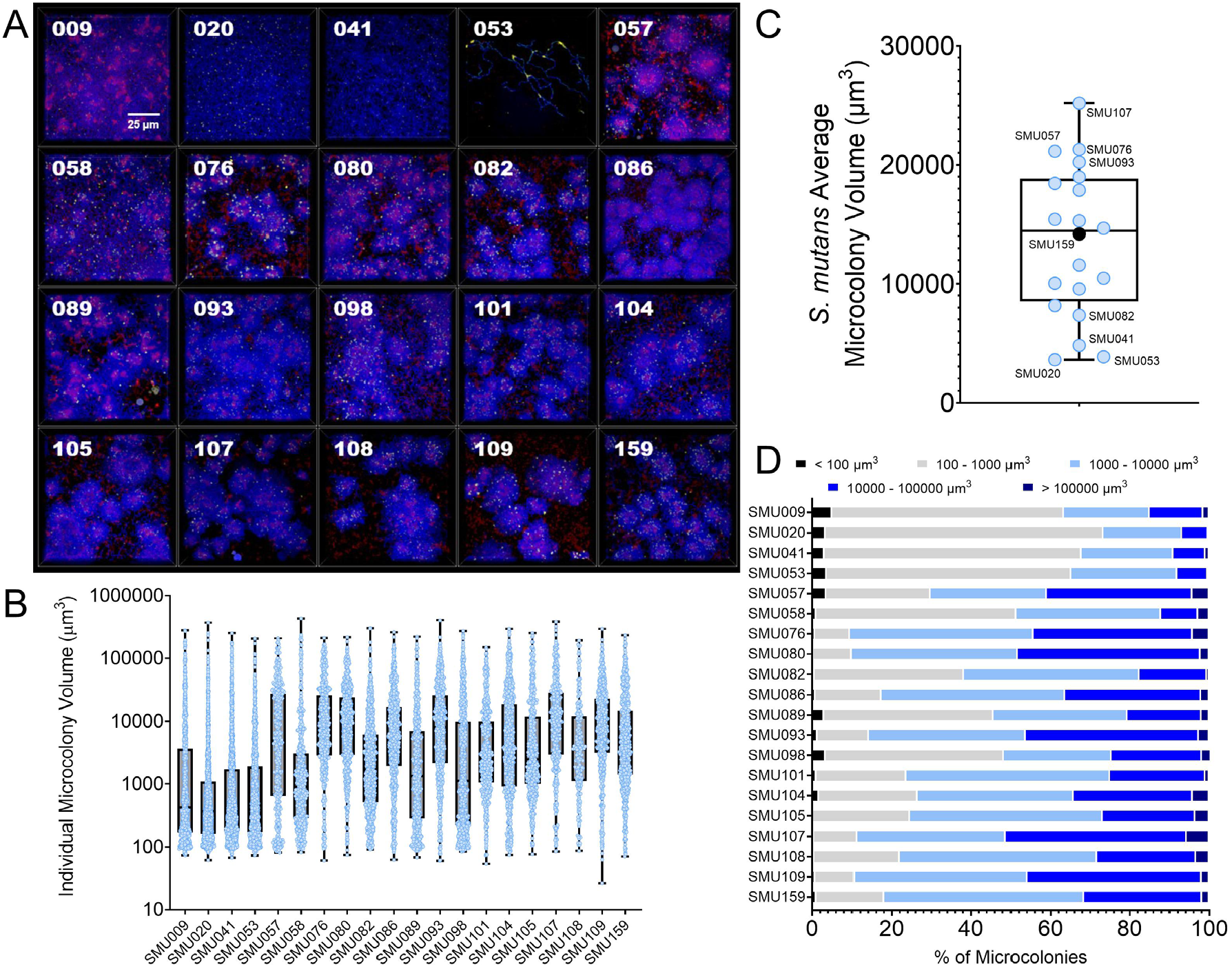
Observation of produced microcolony volumes across a panel of Streptococcus mutans isolates. **(A)** Zoomed-in, maximum intensity 100x 3D merged models of confocal-captured biofilm images oriented top down (Z+) of twenty different *S. mutans* isolates, with the isolate (SMU) number in the top left. Three different channels were captured within each image: total cells (Hoechst total cell stain, blue), eDNA (Alexa Fluor 594-labled α-dsDNA antibody, yellow), and glucans (Alexa Fluor 647-labled dextran, red). Biofilms were grown for 18 h in TYGS medium. Scale bar (25 µm) is shown in the bottom right of the top left merged image. Images are 127 μm (L) x 127 μm (W) x 30 μm (H). **(B)** Microcolony volume of individual microcolonies (blue circles) for each isolate with a box and whiskers plot overlaid capturing the mean and 10^th^ to 90^th^ percentile of volumes. Individual microcolony volumes for the images shown in A were calculated using BiofilmQ. Microcolonies from five separate images (n = 5) were measured and included in the analysis. **(C)** The overall *S. mutans* microcolony volume population average capturing the mean microcolony volume produced by each individual isolate. SMU159 (UA159) is represented by a black dot, while other isolates are labeled. **(D)** Percentage (%) of microcolonies with the indicated volume (figure legend) is shown across all 20 isolates.

When microcolony measurements were averaged by isolate, the panel spanned nearly an order of magnitude **(Figure 1C; Table S1)**. Isolate SMU107 produced the largest average microcolony volume, followed by SMU076 and SMU057, whereas SMU020, SMU053, and SMU041 occupied the lower end of the distribution. UA159 (SMU159) fell near the center of the isolate panel, demonstrating that the commonly used reference strain represents a mid-range architectural phenotype under these conditions. The distribution analysis further distinguished strains that produced predominantly small structures from strains with a large fraction of objects in the 10,000 to 100,000 µm^3^ and greater than 100,000 µm^3^ size classes **(Figure 1D)**. Thus, *S. mutans* microcolony architecture is not a uniform species-level phenotype but instead varies substantially across strain backgrounds. **Supplemental Figure 2** provides full-field views of the merged and individual fluorescence channels, confirming the varieties observed in spatial organization.

### Matrix Accumulation is Associated withss Total Biomass but Does Not Fully Explain Microcolony Architecture

To further characterize biofilm architecture, we quantified the biomass of cells (blue, Hoechst *Strain-level Diversity in Streptococcus mutans* 33342 stain), glucans (red, Alexa Fluor 647-labeled dextran) and extracellular DNA (yellow, α-dsDNA antibody, Alexa Fluor 594-labeled secondary) (11) separately across the panel of *S. mutans* isolate biofilms imaged in Figure 1. **Figure 2A** shows the total cellular biomass for each isolate (left panel) along with a population overview of average biomass for each of the twenty isolates combined (right panel). Total cellular biomass varied across the isolate panel, although the range of average cellular biomass was narrower than the range of individual microcolony volumes (Figure 1). Accumulated glucan biomass displayed pronounced strain-to-strain differences **(Figure 2B)**. SMU020, SMU041, and SMU053 accumulated little detectable glucan and formed few conventional glucan-enmeshed microcolonies. These three strains are known to harbor recombination events in *gtf* genes, which account for their deficient glucan production (16, 18). eDNA biomass also varied, but its isolate-level pattern did not simply mirror glucan accumulation **(Figure 2C)**. To explore relationships between matrix abundance and accumulated cellular biomass, we performed pairwise comparisons of average biomass values across isolates. **Figure 2D** shows association between *S. mutans* accumulated cell biomass and the glucan production by each isolate averaged. Linear regression analysis revealed a weak positive correlation (R^2^ = 0.51), suggesting that higher total cell biomass was associated, but did not always correlate, with increased glucan production. When looking at eDNA accumulation with cell biomass, a similar relationship was observed (R^2^ = 0.54) with some outliers (SMU107 and SMU041) present **(Figure 2E)**. By contrast, the relationship between accumulated glucan and eDNA biomass in the same strain’s biofilm was weaker (R^2^ = 0.35), indicating that isolates differed in the relative accumulation of these two individual matrix components **(Figure 2F)**. Moreover, average microcolony volume was only weakly associated with quantified glucan biomass (R^2^ = 0.27) or eDNA biomass (R^2^ = 0.29) of that same strain **(Figure S3)**. Collectively, these analyses indicate that isolates accumulating more matrix generally amassed more total biofilm biomass, but neither the quantity of glucan nor of eDNA alone could solely determine total biomass or average microcolony size of a given isolate. The data therefore supports a model in which total biomass, matrix composition, and three-dimensional organization are related but separable features of the *S. mutans* biofilm formation phenotype.

**Figure 2.**
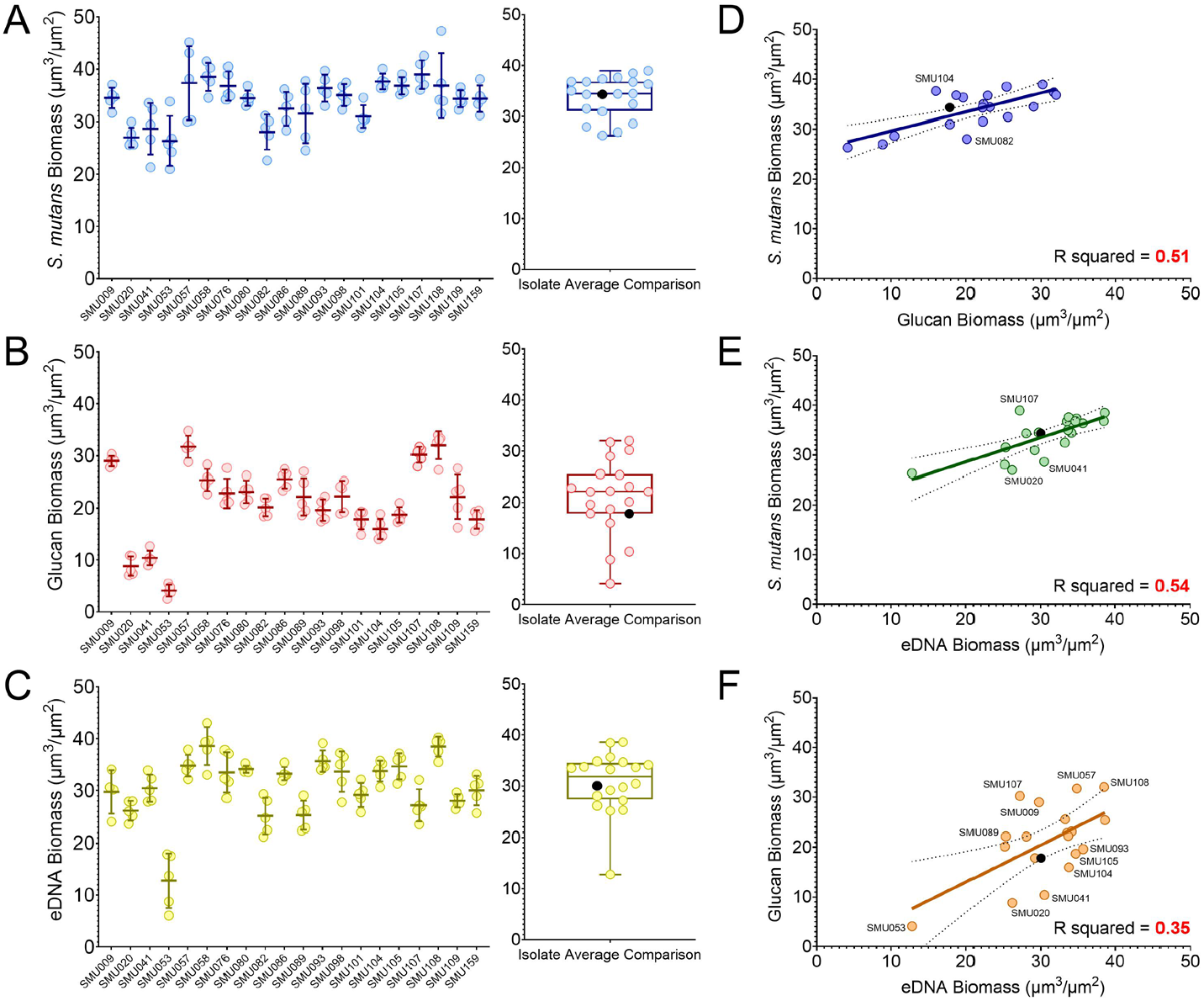
Quantification of extracellular matrix component biomass, and its relation to total cell biomass. Biomass quantification of confocal-captured images shown in Figure 1A, measuring biomass of **(A)** *S. mutans* total cells (blue), along with extracellular matrix components **(B)** glucans (red) and **(C)** eDNA (yellow). Individual strain data (n=5 images analyzed) is plotted on the left while the overall population average (i.e., average for each of the twenty individual isolates shown together) is plotted on the right, with SMU159 (UA159) being represented by a black dot. **(D)** Simple linear regression plot of average *S. mutans* total cell biomass (y-axis) by average glucan biomass of that same isolate (x-axis). Averages were taken from data shown in panels A-C. SMU159 is represented by a black dot, while other outlier strains are labeled. **(E)** Simple linear regression plot of average *S. mutans* total cell biomass (y-axis) by average eDNA biomass of that same isolate (x-axis). **(F)** Simple linear regression plot of average glucan biomass (y-axis) by average eDNA biomass of that same isolate (x-axis). Linear regression and R squared values were calculated using built-in analysis within GraphPad Prism (v11.0). Calculated R squared values are shown in bottom right corner in bolded red.

### Human Saliva Produces Strain-Specific Remodeling of Biofilm Architecture and Matrix Composition

Previously, our group reported that addition of human saliva to the culture media alters biofilm formation by several oral streptococci, including *S. mutans* (19). To determine whether this environmental signaling extends across a broader set of isolates, we evaluated biofilm formation in TYGS with or without saliva (1:1, v/v).

Representative confocal images from eight isolates **(Figure 3A)** demonstrate that presence of saliva leads to marked architectural changes in *S. mutans* isolate biofilms, with visible changes in microcolony spatial organization.

**Figure 3.**
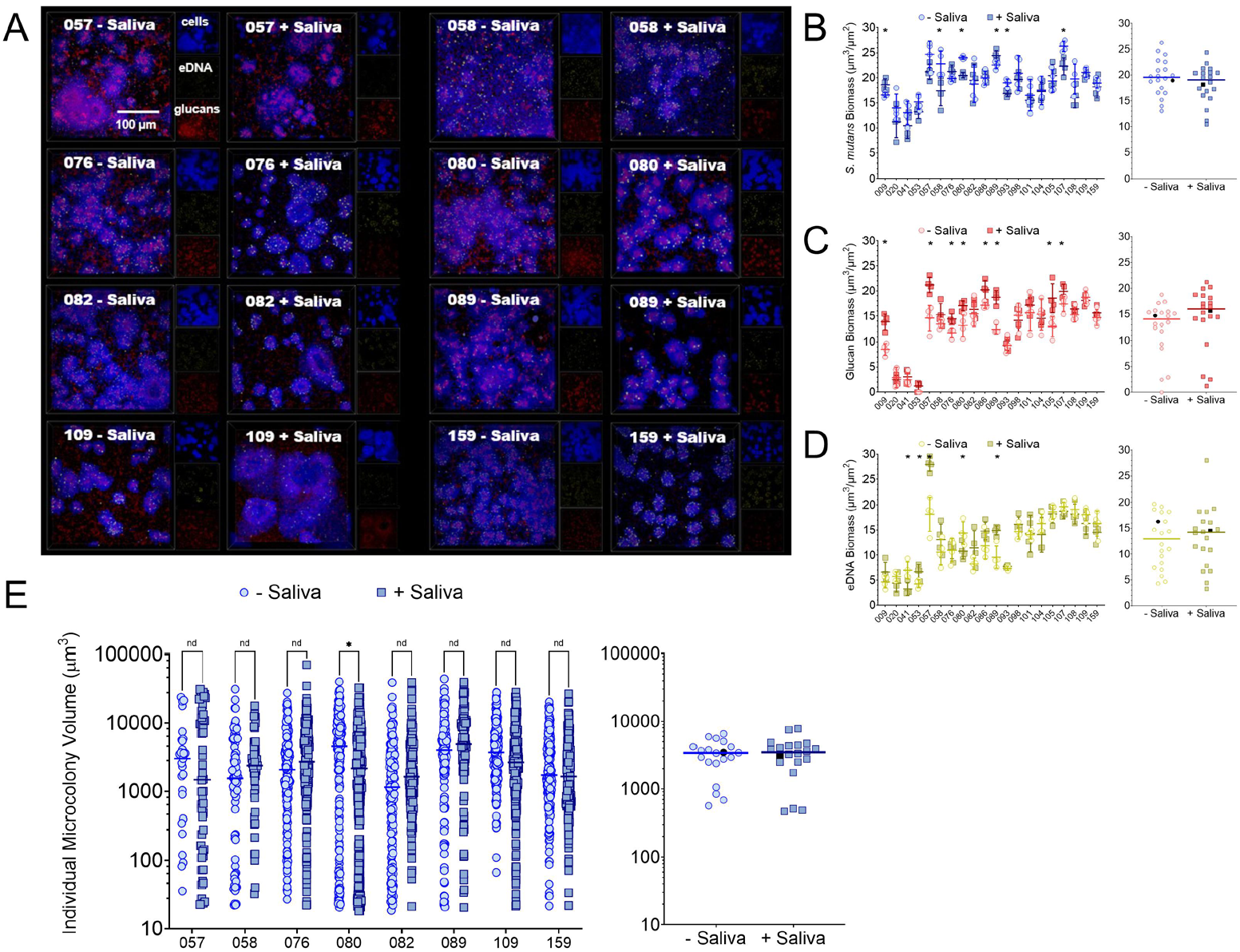
Impact of human saliva on the biofilm architecture of Streptococcus mutans microcolony formation. **(A)** Maximum intensity, 100x 3D merged models of confocal-captured biofilm images oriented top down (Z+) of eight different *S. mutans* isolates, with the isolate (SMU) number and condition listed on the top of the image. Three different channels were captured within each image: total cells (blue), eDNA (yellow), and glucans (red). Biofilms were grown for 18 h in TYGS medium either with (+ Saliva) or without (-Saliva) human saliva (1:1, v/v). Images of individual channels showing accumulation and spatial distribution of each component are shown on the right of the merged image. Scale bar (100 µm) is shown in the bottom right of the top left merged image. Images are 256 μm (L) x 256 μm (W) x 30 μm (H). Biomass quantification of captured images shown in Figure 3A, measuring biomass of **(B)** *S. mutans* total cells (blue), along with extracellular matrix components **(C)** glucans (red) and **(D)** eDNA (yellow) in TYGS medium either with (+ Saliva, squares, darker color) or without (-Saliva, circles, lighter color) saliva. Individual strain data (n=5 images analyzed) is plotted on the left while the overall population average (i.e., average for each of the twenty individual isolates shown together) is plotted on the right, with SMU159 (UA159) represented by a black circle/square. Line shown represents the mean biomass for the overall population. **(E)** Microcolony volume of individual microcolonies for each isolate shown in 3A, calculated using BiofilmQ. Microcolonies from three separate images were measured and included in the analysis. Multiple unpaired t-tests with Welch correction were conducted between -saliva and +saliva conditions, * = P value < 0.01, nd = no significant difference.

At the isolate level, saliva altered total cell biomass in several strains, including SMU009, SMU058, SMU080, SMU089, SMU093, and SMU107 **(Figure 3B)**.

The overall tendency was toward reduced cellular biomass in saliva-containing medium, but this response was not universal, and aggregation of isolate averages did not reveal a significant population-wide shift. Matrix components showed a similarly heterogeneous response. Glucan biomass increased in several isolates, including SMU009, SMU057, SMU076, SMU080, SMU086, SMU089, SMU105, and SMU107 **(Figure 3C)**, whereas significant eDNA changes were observed for a smaller subset that included SMU053, SMU057, and SMU093 **(Figure 3D)**. Despite these isolate-specific effects, the population-wide isolate averages for glucan and eDNA were not significantly different between the two conditions.

Analysis of individual microcolony volumes reinforced the conclusion that saliva responsiveness was strain dependent **(Figure 3E)**. Among the eight isolates selected for detailed microcolony analysis, only SMU080 showed a significant shift in microcolony volume distribution with saliva, even though visible architectural changes were apparent in several image panels. The absence of a uniform population-level response, together with distinct isolate-specific changes in cells and matrix, indicates that saliva acts as an environmental modifier whose effect depends on strain background.

### S. mutans Isolates Differ in Growth and Retained Biomass Under Acidic Starting Conditions

To determine whether strain-level microcolony architectural diversity was accompanied by differences in acid-stress performance, we evaluated planktonic growth and attached biomass after inoculation of each isolate into TYG(S) media pre-adjusted to starting pH values of 7.0, 6.0, or 5.5 (20). Growth patterns were quantified by measuring doubling time, final yield (24 h), lag phase duration (min to OD_600_ = 0.1) and CV staining (24 h). Individual growth curves of each isolate used to generate the heatmaps are shown in **Figure S4**.

Across the panel, decreasing the initial pH generally increased doubling time, decreased final yield, and prolonged the time required to reach an OD600 of 0.1 **(Figures 4A-C)**. However, the magnitude of these effects varied substantially among isolates. At pH 7.0, most isolates entered exponential growth within a similar general time frame, although baseline growth rates and final yields were not identical. At pH 6.0 and particularly at pH 5.5, several strains showed large increases in doubling time and pronounced reductions in yield, whereas others retained comparatively stronger growth such as UA159. SMU107 was a notable isolate as its lag-associated phenotype was altered less than most other isolates in the panel as the starting pH decreased. The heatmaps therefore revealed multiple dimensions of low-pH performance: an isolate could retain a relatively short lag yet still exhibit a reduced final yield or maintain biomass while growing more slowly.

**Figure 4.**
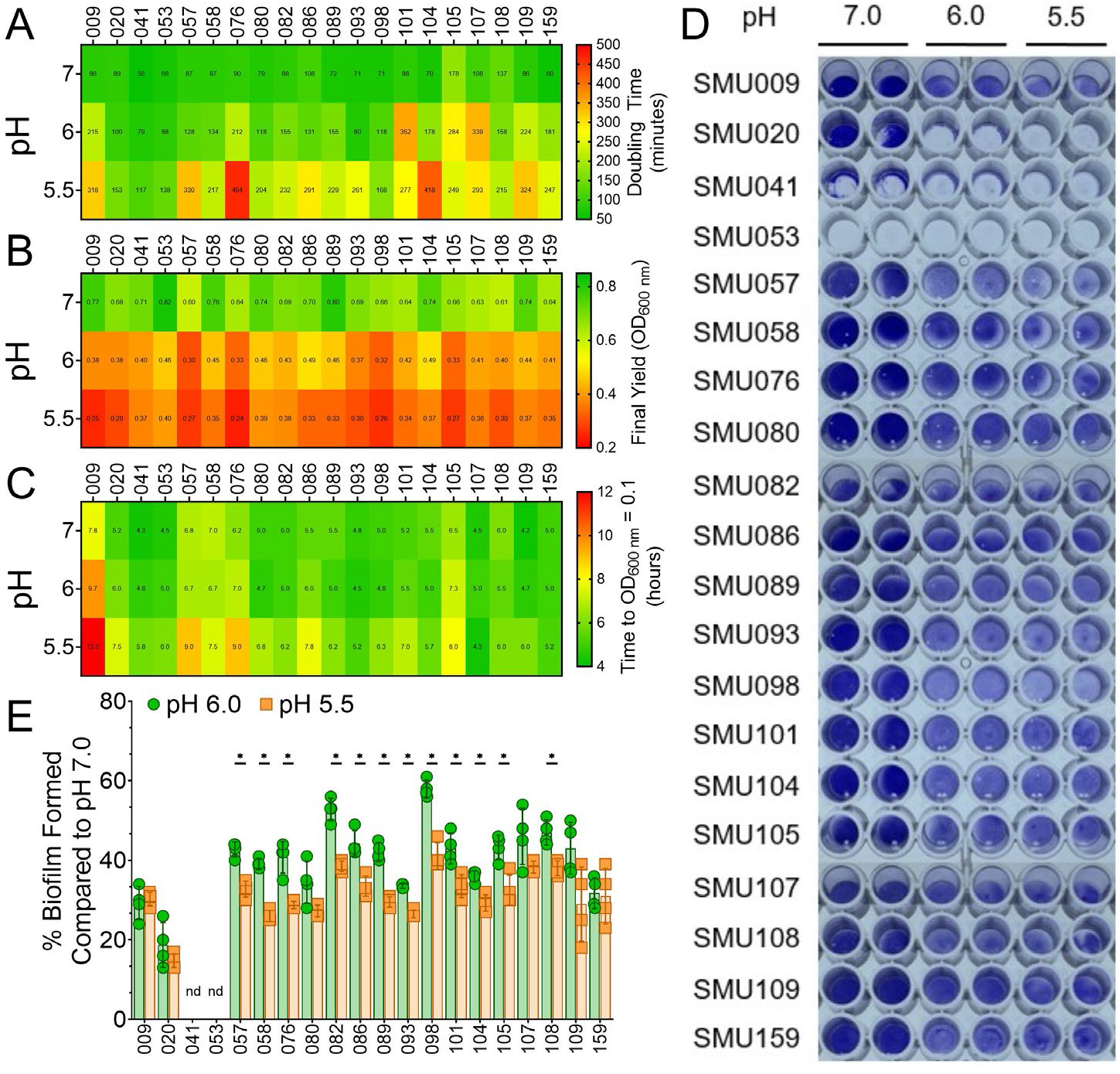
Impacts of initial pH on growth and biofilm formation for a panel of Streptococcus mutans isolates. Heatmaps of *S. mutans* isolate **(A)** doubling times in minutes, **(B)** final yields after 24 h of growth (recorded optical density at 600 nm), and **(C)** time in hours to surpass OD600 nm = 0.1 (i.e., lag time). Strains were grown in TYG medium with initial pH of the media adjusted to either pH 7.0, 6.0 or 5.5. Growth was monitored within a Bioscreen C for 24 h with a reading every 0.5 h. The highest values within the heatmap are shown in green, intermediate values in yellow, while the lowest values are in red. Actual recorded mean values are shown within each cell of the heatmap. **(D)** Representative images of a CV biofilm biomass assay where individual *S. mutans* isolates were cultured in TYGS medium with initial pH of the media adjusted to either pH 7.0, 6.0 or 5.5. Individual *S. mutans* isolates are listed down the left y-axis in numerical order from lowest to highest, while initial pH values of the media are listed on the top x-axis. **(E)** Quantification of the CV assay shown in Figure 4D, expressed as percentage biofilm formed (OD575 nm values) compared to the pH 7.0 condition. Percentage biofilm formed in pH 6.0 medium is shown as light green circles, while pH 5.5 is shown in dark orange squares. Multiple unpaired t-tests with Welch correction was conducted between pH 6.0 and pH 5.5, * = P value < 0.01. n = 4, nd = values not determined due to low initial biofilm formation at pH 7.0 (SMU041 and SMU053).

We also quantified CV-stained biomass retention after growth at the three starting pH values **(Figure 4D and 4E)**. Because isolates differed in their baseline attachment at pH 7.0 (i.e., differed in OD_575 nm_ values), biomass at pH 6.0 and pH 5.5 were expressed relative to the pH 7.0 value for each strain. Most isolates retained less attached biomass as the starting pH decreased, but the degree of reduction varied. SMU076, SMU080, SMU086, SMU107, SMU108, and SMU109 retained comparatively high proportions of their pH 7.0 biomass at pH 5.5, whereas SMU020 showed a pronounced reduction. SMU041 and SMU053 could not be evaluated in this normalized analysis because their baseline biomass at pH 7.0 was too low for a meaningful comparison (not determined, nd).

These results demonstrate that planktonic growth rate, lag time, final yield, and attached biomass respond differently across isolates when cultures are initiated at low pH. Accordingly, acid tolerance performance cannot be represented by a single measurement, and the isolates display distinct combinations of planktonic growth and biofilm-associated phenotypes.

### Relative Acid Accumulation Varies Among Isolates Independent of Microcolony Volume

We next evaluated whether isolates that formed larger microcolonies also generated or retained more acid within mature biofilms. To measure this, 18 h TYGS biofilms were stained with pHrodo Green AM (a pH-sensitive dye that produces higher fluorescent intensity in more acidic environments) together with a total cell stain (Hoechst 33342). In a validation experiment using GFP-expressing *S. mutans* UA159 (21, 22) cocultured with *Streptococcus gordonii* (distinguished by the total cell stain only), pHrodo fluorescence localized predominantly to *S. mutans* microcolonies rather than the surrounding commensal population **(Figure S5A)**. In that experimental context, fluorescence increased with individual microcolony volume in a nonlinear relationship (R^2^ = 0.68; **Figure S5B**), supporting the use of the dye as a relative indicator of acid-associated microenvironments.

Images of the pHrodo channel alone for the full isolate panel are shown in **Figure S6**. Across the strains, pHrodo fluorescence varied widely **(Figures 5A and 5B)**. SMU058, SMU076, SMU105, and SMU107 were among the isolates with a strong average fluorescent signal, whereas SMU009, SMU098, SMU101, SMU108, SMU109, and UA159 displayed lower average signal under the conditions tested. Considerable heterogeneity was also present among individual microcolonies produced by the same isolate, showing that acid-associated fluorescence was not spatially uniform within a single strain’s population.

**Figure 5.**
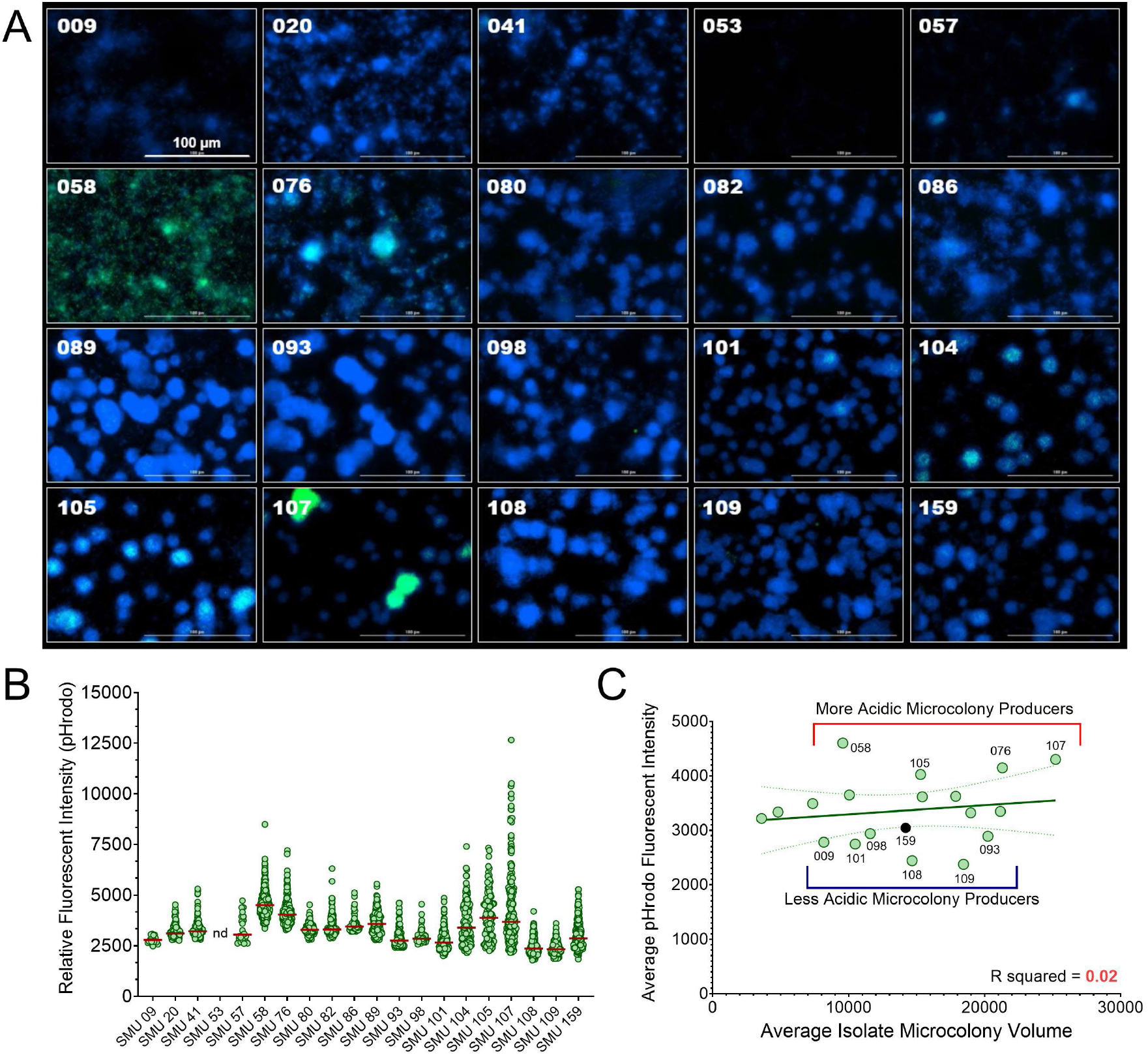
Acid-Associated Microcolony Fluorescence Varies Across Streptococcus mutans Isolates Independently of Microcolony Volume. **(A)** Merged representative maximum intensity 40x Z projection images of 18 h *S. mutans* biofilms, with a total cell stain shown in blue (Hoechst 33342), and pHrodo Green AM pH indicator shown in green. The isolate SMU number for each representative image is shown in the top left. Scale bar (100 µm) is shown in the bottom right of the top left image. Images are 256 μm (L) x 256 μm (W) x 30 μm (H). **(B)** Relative fluorescent intensity of the pH indicator dye within individual microcolonies from the images shown in 5A. Each data point represents a recorded fluorescent intensity from an individual microcolony, while the red line represents the mean fluorescent intensity for that particular isolate. Fluorescent intensities were calculated using Gen5 Image+ software (v 3.11). Microcolonies from five separate images (n = 5) were measured and included in the analysis. **(C)** Simple linear regression plot of average *S. mutans* isolate pH indicator dye fluorescent intensity (y-axis) by average microcolony volume of that same isolate (x-axis). Linear regression and R squared values were calculated using built-in analysis within GraphPad Prism (v11.0). Calculated R squared value is shown in bottom right corner.

At the isolate level, average pHrodo fluorescence was not associated with average microcolony volume (R^2^ = 0.02; **Figure 5C**). Thus, strains that produced the largest structures were not necessarily those with the strongest acid-accumulation signal. The comparison with low-pH growth phenotypes also revealed discordant trait combinations. For example, SMU107 combined large microcolonies, strong pHrodo fluorescence, and comparatively robust performance under acidic starting conditions making it an isolate with high cariogenic potential that warrants further investigation. In contrast, SMU076 produced strong pHrodo fluorescence despite more pronounced impairment in some planktonic growth measurements at low initial pH. These findings distinguish relative acid accumulation within a mature biofilm from the ability to initiate and sustain growth under acidic conditions and demonstrate that both properties vary independently of average microcolony volume across the isolate panel.

### Oral Health Antimicrobials Inhibit Biofilm Accumulation in Isolate- and Compound-Specific Patterns

Mature biofilms are thought to limit antimicrobial penetration, potentially conferring increased tolerance to antimicrobial agents (6, 23). We therefore wanted to examine whether the ability of specific *S. mutans* isolates to form larger microcolony structures influenced their susceptibility to two commonly used cationic antiseptics in oral healthcare products and clinical practice (24, 25). To address this, we evaluated the susceptibility of *S. mutans* isolate panel to chlorhexidine (CHX) and cetylpyridinium chloride (CPC).

First, the minimal biofilm inhibitory concentration (MBIC) was evaluated for each isolate to either CHX or CPC **(Figure 6A and 6B, respectively)**. The CV-stained biofilms (right) and corresponding quantitative measurements (left) are shown, with biofilm biomass expressed as a % relative to untreated controls (i.e., 0.00 μg mL^-1^ CHX or CPC). For CHX, most isolates exhibited a sharp inhibitory threshold between 0.75 and 1.0 µg mL^-1^, while certain isolates, such as SMU020 showed minimal biofilm formation and thus, a higher loss of biomass at lower concentrations (0.25 µg mL^-1^). Notably, SMU076 and SMU082 showed tolerance to higher concentrations of CHX, maintaining 98% and 74% of biomass, respectively, even at 0.75 µg mL^-1^. In comparison, CPC treatment resulted in a broader inhibitory profile. Many isolates maintained substantial biomass at 0.50 – 1.0 µg mL^-1^ CPC, with inhibition becoming evident only at ≥2.0 µg mL^-1^. SMU020 again displayed low tolerance to CPC, consistent with its response to CHX. In contrast, SMU107 showed stronger inhibition by CPC than would have been predicted from its CHX response. Isolates SMU086 and SMU093 were more tolerant to CPC but much more sensitive to CHX. Notably, SMU076 was among the most tolerant isolates to both CHX and CPC, maintaining high biomass across a range of concentrations. The antiseptic susceptibility patterns did not recapitulate the ranking of an isolate’s average microcolony volume (Table S1). Isolates that produce large microcolonies were not uniformly the most tolerant to the antiseptics, and isolates that produce smaller microcolonies were not uniformly the most susceptible. These experiments therefore show that inhibition of biofilm accumulation is strongly influenced by both strain background and antimicrobial compound, and cannot be predicted from displayed biofilm spatial organization alone.

**Figure 6.**
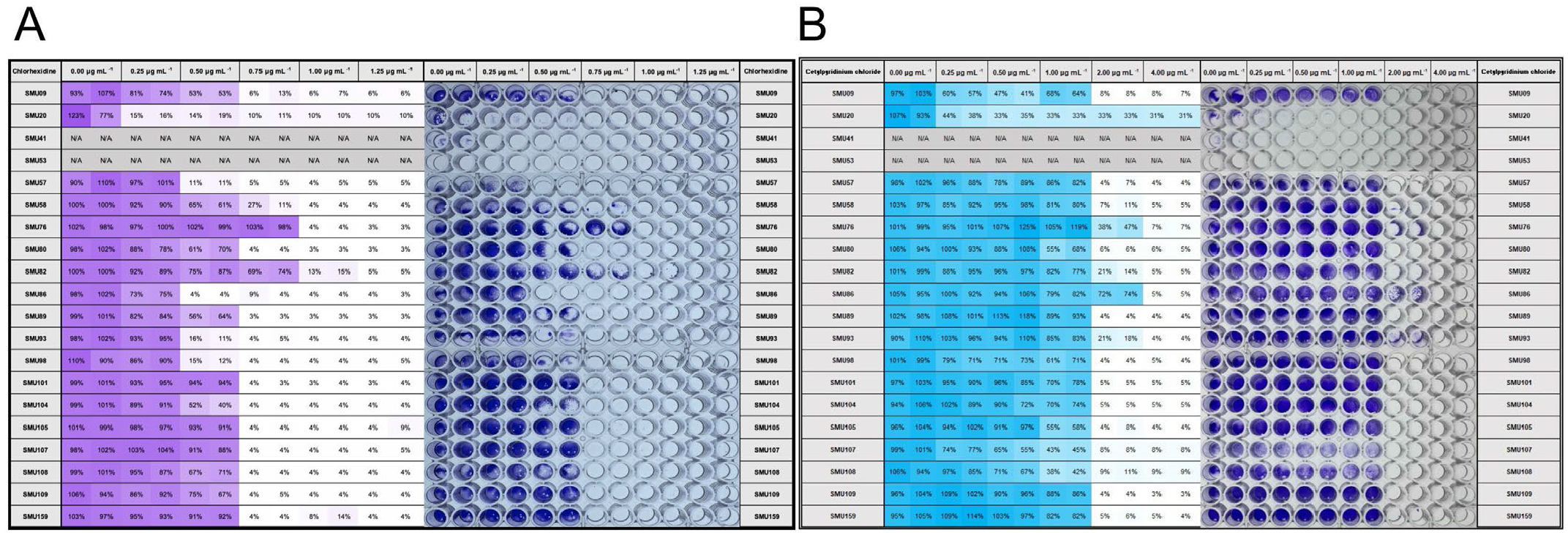
Minimal biofilm inhibitory concentration (MBIC) of a panel of Streptococcus mutans isolates against antiseptics chlorhexidine and cetylpyridinium chloride. Representative images of a CV biofilm biomass assay where individual *S. mutans* isolates were cultured in TYGS medium with increasing concentrations of either **(A)** chlorhexidine (CHX) or **(B)** cetylpyridinium chloride (CPC). Individual *S. mutans* isolates are listed down the left y-axis in numerical order from lowest to highest, while CHX and CPC concentrations (µg mL^-1^) are listed on the top x-axis. A quantification of the CV is shown to the left of the image, expressed as percentage biofilm formed (OD 575 nm) compared to no antimicrobial added (0.00 µg mL^-1^). The highest values are shown as dark purple or blue, while the lowest values are white. Actual recorded values are shown within each cell. N/A = not available, values not determined due to low initial biofilm formation at 0.00 µg mL^-1^ (SMU041 and SMU053).

### Brief Antiseptic Exposure Reveals Isolate-Specific Baseline Membrane Permeability and a Strong Dose Response

Because continuous exposure during growth does not effectively model the brief contact time of a biofilm during an oral rinse of a mouthwash product, we separately examined established 18 h biofilms after a thirty second CHX challenge. Seven isolates representing different architectural and susceptibility phenotypes were treated with either 50 or 500 µg mL^-1^ CHX and later stained with a total cell stain (Hoechst 33342) and SYTOX Green, which assesses cell membrane integrity. PBS and ethanol served as negative and positive controls, respectively **(Figure 7A)**. The untreated biofilms differed substantially in baseline SYTOX positivity **(Figure 7B)**. SMU082 displayed a comparatively high fraction of membrane-compromised cells in the PBS control condition, whereas SMU076 and SMU107 showed low baseline SYTOX staining. Following exposure to 50 µg mL^-1^ CHX, the response remained isolate dependent. Among the tested strains, SMU107 was the only isolate that showed a significant increase relative to its PBS control. In contrast, 500 µg mL^-1^ CHX produced high SYTOX fluorescence across all seven isolates, approaching the ethanol-treated positive controls.

**Figure 7.**
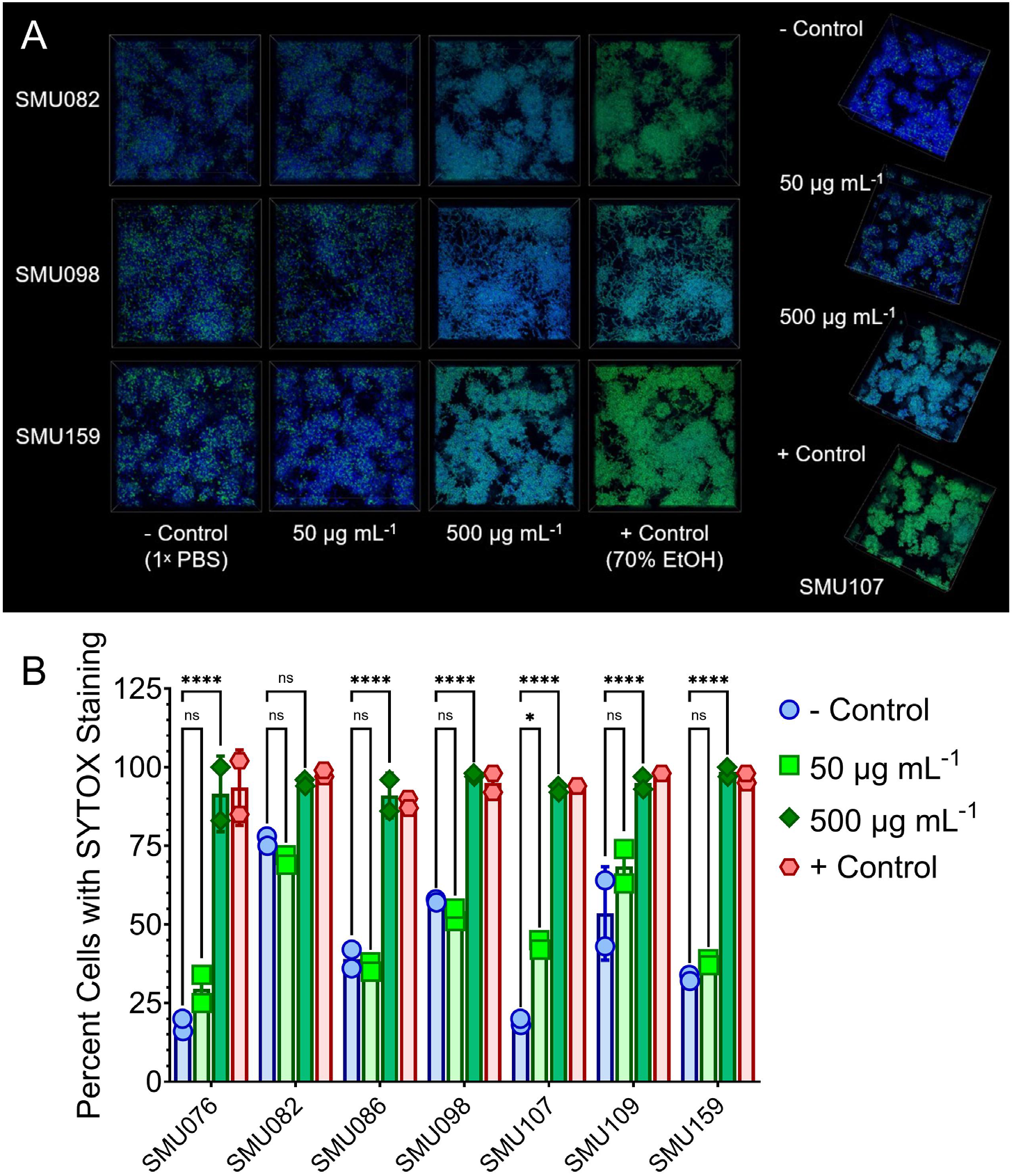
Impacts of short term exposure of Streptococcus mutans isolates to chlorhexidine. **(A)** Maximum intensity, 100x 3D merged models of confocal-captured biofilm images oriented top down (Z+) of four different *S. mutans* isolates, with the isolate (SMU) number listed, and condition to the bottom or to the side. Biofilms were grown for 18 h in TYGS medium. After 18 h, biofilms were treated with either 50 or 500 µg mL^-1^ chlorhexidine (CHX) for thirty seconds, or with 1× PBS (negative control) or 70% ethanol (EtOH; positive control). Biofilms were then stained with a total cell stain shown in blue (Hoechst 33342) along with SYTOX Green (green), which is impermeable to the cell unless the cell membrane is compromised. Images are 256 μm (L) x 256 μm (W) x 30 μm (H). **(B)** Quantification of the images shown in Figure 7A, expressed as percentage cells with SYTOX staining, indicating compromised membranes with their respective treatments. Seven total isolates were used in the analysis, with percentages determined for the negative control (-Control; blue circles), addition of 50 µg mL^-1^ CHX (medium green squares), addition of 500 µg mL^-1^ CHX (dark green diamonds), and the positive control (+ Control; red hexagons). Statistics were completed with an ordinary two-way ANOVA which included Tukey’s multiple comparisons test, * = P value < 0.05, **** = P value < 0.0001, ns = not significant.

Qualitative inspection of the confocal images did not reveal an obvious spatial pattern in which SYTOX-positive cells were restricted to microcolony surfaces or consistently excluded from microcolony interiors. Therefore, within the spatial resolution and treatment conditions used here, large microcolonies did not provide an evident zone of complete protection from membrane damage at higher CHX concentrations. Together with the continuous-exposure assays, these data indicate that baseline membrane integrity after biofilm formation, inhibition of biofilm accumulation, and the response of established biofilms to a brief antiseptic challenge are related but distinct strain-level phenotypes.

## DISCUSSION

In this study, we revisited a previously characterized collection of genomically and phenotypically diverse *Streptococcus mutans* isolates (16) to examine strain-level biofilm variation in microcolony architecture, extracellular matrix composition, and functional phenotypes relevant to cariogenicity. Using quantitative super-resolution confocal microscopy together with assays of acid-associated physiology and antimicrobial susceptibility, we identified substantial differences among isolates in biofilm organization, matrix accumulation, acid accumulation and tolerance, and responses to oral antiseptics. These findings demonstrate that disease-relevant traits commonly attributed to *S. mutans* at the species level are distributed unevenly across genetically diverse strains. Importantly, these phenotypes did not consistently vary together as a single coordinated program. Instead, our results support a model in which the pathogenic potential of *S. mutans* emerges from partially coupled functional trait modules rather than from any one dominant phenotype, such as displayed average microcolony volume.

Our data demonstrates substantial heterogeneity in how *S. mutans* isolates form microcolony structures (Figure 1 and 2). Individual isolates generated distributions of microcolony sizes spanning several orders of magnitude, and isolate-average volumes varied nearly tenfold. UA159 occupied the middle of this distribution, reinforcing its value as a well-characterized experimental reference while also showing the limitation of treating one strain as representative of the species. The isolates also varied widely in the relative contribution of extracellular matrix components, including polysaccharide glucans and extracellular DNA (eDNA) (Figure 2). While modest trends were observed between total biomass accumulation and EPS production, these patterns were weaker when considering average microcolony volume, suggesting that EPS abundance alone does not dictate microcolony size or architecture. Notably, some strains preferentially accumulated eDNA over glucans, whereas others displayed the opposite pattern, highlighting the diversity of matrix composition strategies employed by different isolates.

Interestingly, microcolony architecture did not predict acid accumulation within biofilms. Although acid production is a central virulence trait of *S. mutans*, we observed no correlation between average microcolony volume and the degree of microcolony acidification at the 24-hour time point. This finding suggests that acidogenicity is governed by factors independent of biofilm volume or structure. Instead, genetically encoded factors, such as accessory genes, differences in regulatory gene networks or variability in metabolic enzyme activity, are likely to play a role in determining acid production at the individual strain level (26–29). Despite this lack of correlation, we identified a subset of isolates, such as SMU058 and SMU107, that consistently formed highly acidic microcolonies despite occupying distinct positions on the phylogenetic tree (Figure S1). These strains may represent a functionally distinct group with enhanced acidogenic potential, independent of their produced biofilm architectural features. A limitation of the present study is the absence of detailed clinical metadata associated with these isolates, preventing direct correlation between observed phenotypes and disease status or lesion activity at the time of isolation to understand clinical implication. A future study of interest may be to isolate new strains of *S. mutans* with known clinical parameters and determine if isolates from more progressed disease or lesion site show similar highly acidogenic phenotypes.

Our findings further indicate that biofilm architecture and microcolony formation do not confer predictable tolerance to commonly used oral healthcare antiseptics. Susceptibility to chlorhexidine (CHX) and cetylpyridinium chloride (CPC) varied markedly across isolates and differed depending on the antimicrobial tested (Figure 6). Neither microcolony size nor biofilm structure correlated with MBIC values or cell death following short-term CHX exposure. Instead, antimicrobial tolerance displayed isolate-specific physiological traits, likely including differences in membrane properties, stress response pathways, or detoxification mechanisms, rather than physical properties from the biofilm. An intriguing observation was that some isolates showed more permeable membranes after 24 hours than others, even in the control condition (i.e., no antiseptic exposure). For example, SMU082 displayed substantial SYTOX staining at baseline, whereas SMU107 showed very low membrane permeability (Figure 7B). Of note, SMU107 demonstrated high acid tolerance, with minimal changes in growth rate and sustained high biomass accumulation even at pH 5.5 (Figure 4). However, this acid tolerance characteristic was not consistently associated with antimicrobial susceptibility profiles across isolates, suggesting that these phenotypes are governed by partially distinct mechanisms.

These results reflect isolate-specific variations in cell envelope integrity and composition, including differences in membrane lipid profiles, cell wall thickness or teichoic acid modification, all of which can influence membrane permeability, stress tolerance, survival under acidic or antimicrobial challenge (30–32). Additionally, differences in autolytic activity may further contribute to the observed variability in membrane permeability and cell death. The autolysin AtlA, which is a significant contributor to *S. mutans* autolysis (33, 34), has previously been shown to vary in activity across these *S. mutans* clinical isolates (16), with some strains showing reduced autolysis compared to UA159 and others displaying minimal autolytic activity. These observations suggest that baseline differences in membrane permeability, acid tolerance and antimicrobial response among *S. mutans* isolates arise from complex, strain-specific variation in cell envelope physiology.

Our work highlights not only the heterogeneity of *S. mutans* isolates but also the complexity of *S. mutans* pathogenicity and underpins the notion that cariogenic potential is not derived from a single dominant phenotypic trait. Instead, key disease-relevant properties, such as biofilm architecture, acidogenicity, aciduricity and antimicrobial tolerance can differ substantially across strains and are not part of a single coordinated virulence axis. In this context, individual strains that colonize with complementary phenotypes may collectively enhance biofilm persistence, acid accumulation and stress tolerance within the microbial community and promote disease progression. As the patient-level disease status for clinical isolates examined in this study is incomplete, it remains unclear whether particular phenotypes, i.e. high acidogenicity or robust biofilm formation, are enhanced in isolates obtained from caries-active individuals as an evolutionary consequence. Alternatively, these findings raise the possibility that substantial phenotypic diversity exists even within a single host. Within a caries-active individual, distinct *S. mutans* strains may occupy complementary ecological niches, with some strains contributing disproportionately to acid production while others provide structural stability through enhanced biofilm formation or stress tolerance (35–40). Such functional specialization could collectively enhance biofilm persistence and acid retention at the tooth surface, thereby driving disease progression. The current experiments do not demonstrate cooperation among strains, but they raise a testable ecological hypothesis: strain-level functional diversity could broaden the collective capabilities of an *S. mutans* population within a polymicrobial biofilm. The heterogeneity in various disease-relevant phenotypes we observed in *S. mutans* isolates also underscores the limitations of species-level detection or quantification of *S. mutans* as an indicator for caries risk (41, 42). The presence of *S. mutans* alone therefore may be insufficient to predict disease activity, highlighting the need for ecology-informed, precision-based therapeutic approaches that account for strain-level diversity and cooperative behaviors within oral biofilms.

In summary, *S. mutans* isolates differ not only in the magnitude of individual cariogenic-associated traits but also in the combinations in which those traits occur. The widely used reference strain UA159 displays a useful intermediate phenotype but does not capture the structural, physiological, or antimicrobial-response extremes observed across the species. The absence of a consistent relationship among biofilm spatial organization, acid accumulation, acid-stress performance, and antiseptic response cautions against inferring functional behavior from simple species abundances or from any single biofilm measurement. Although not examined here, interactions with commensal streptococci, other oral genera such as *Actinomyces* and *Rothia*, or cross-kingdom partners such as *Candida albicans* may further modify or amplify strain-specific phenotypes within complex plaque communities. Developing a strain-resolved and multidimensional understanding of *S. mutans* biology will therefore be important for defining the functional organization of cariogenic biofilms and for designing interventions that remain effective across the phenotypic diversity present within the species.

## MATERIALS AND METHODS

### Bacterial Strains and Growth Conditions

The *S. mutans* isolates used in this study are listed in **Table S1** and were selected from a previously characterized collection (16). *S. mutans* UA159 (SMU159) was included as a reference strain. All suppliers and catalog numbers for materials used in this study are listed in **Table S2**. Bacterial strains were maintained for long-term storage at −80°C in Brain Heart Infusion (BHI) medium containing 25% glycerol. Overnight cultures were initiated from single, isolated colonies grown on Brain Heart Infusion agar, inoculated into BHI broth, and incubated at 37°C in 5% CO_2_.

The following day, overnight cultures were harvested by centrifugation, washed to remove residual BHI medium, and normalized to an OD_600_ of 0.2 in 1× phosphate-buffered saline (PBS) prior to inoculation into experimental media. Unless otherwise indicated, strains were back diluted 1:100 into tryptone-yeast extract medium (TY; 10 g tryptone, 5 g yeast extract, and 3 g K_2_HPO_4_ per L H_2_O) supplemented with 10 mM glucose (TYG). For all biofilm experiments, 5 mM sucrose was additionally included (TYGS). Cultures were incubated at 37°C in 5% CO_2_ unless otherwise indicated.

### Human Saliva Preparation

Commercially available pooled human saliva was used (19). Upon receipt, saliva was thawed, centrifuged at 4500 rpm for 10 min, and passed through a 0.22-µm filter unit. Processed saliva was divided into 10-mL aliquots and stored frozen until use. For experimental assays, aliquots were thawed and used the same day. Saliva-containing medium was prepared by mixing TY medium and human saliva at a 1:1 ratio (v/v), followed by supplementation with glucose and/or sucrose.

### Biofilm Growth and Extracellular Matrix Labeling for Microscopy

For visualization of biofilm architecture and extracellular matrix components, bacterial strains were inoculated into TYGS containing 1 µM Alexa Fluor 647-labeled dextran (10,000 molecular weight; anionic, fixable), which was incorporated during biofilm formation/glucan synthesis. Cultures were inoculated into 12-well glass-bottom black plates and incubated at 37°C in 5% CO_2_ for 18 h (22).

Following incubation, biofilms were gently washed with 1× PBS to remove loosely associated cells and incubated for 30 min at room temperature in blocking buffer consisting of 3% bovine serum albumin (BSA) in PBS. Extracellular DNA (eDNA) was labeled by incubation with a murine monoclonal antibody recognizing dsDNA (anti-dsDNA, 35I9 DNA) at 2 µg mL^-1^ in BSA blocking buffer for 1 h at room temperature. Biofilms were washed and subsequently incubated for 1 h at room temperature with Alexa Fluor 594-labeled, highly cross-absorbed goat anti-mouse IgG secondary antibody at 2 µg mL^-1^. Following an additional wash, total bacterial cells were stained with Hoechst 33342 at a final concentration of 5 µM for 15 min. Biofilms were maintained in 1× PBS during imaging.

### Super-Resolution Confocal Microscopy and Image Analysis

Super-resolution confocal imaging was performed using a Nikon A1R confocal microscope. Biofilms were visualized using a 100× oil-immersion objective (1.45 numerical aperture; 0.13-mm working distance) with fluorescence collected in the appropriate channels for Hoechst 33342/DAPI, Alexa Fluor 594, and Alexa Fluor 647. A resonant scanner was used for image acquisition, and z-stacks were collected over a field measuring approximately 127.3 × 127.3 µm with a total z-depth of 30 µm and 0.5-µm step intervals. Image acquisition and processing were performed using NIS-Elements software (v 6.02.03). The NIS-Elements Denoise.ai function was applied uniformly across image sets prior to subsequent visualization and quantitative analysis. Maximum-intensity projections and three-dimensional reconstructions were generated from the processed z-stacks. Microcolony volumes were calculated by importing single-channel images corresponding to the total-cell signal into BiofilmQ (43). A fluorescence-intensity threshold greater than 2,000 arbitrary units (a.u.) and a minimum object size greater than 5 µm were applied uniformly during microcolony segmentation. Objects contacting the image boundary were excluded. Five independent image fields were analyzed for each isolate for the primary microcolony analysis in Figure 1. Individual microcolony volumes were exported for subsequent statistical analysis and were also averaged to generate isolate-level mean values. For the saliva comparison, microcolony volumes were determined from three image fields for each of the eight isolates selected for detailed architectural analysis.

Total cellular, glucan, and eDNA biomass were quantified separately from their respective fluorescence channels using BiofilmQ. A common set of segmentation and intensity parameters was maintained across isolates within each fluorescence channel to permit direct comparison among strain backgrounds. Five image fields were analyzed for each isolate. Biomass values from individual image fields were averaged to generate isolate-level values used for correlation analyses.

### Crystal Violet Biofilm Biomass Quantification

Biofilm biomass was quantified using a crystal violet (CV) retention assay as previously described (44). Strains were inoculated into 96-well plates containing the indicated experimental medium and incubated for 24 h at 37°C in 5% CO_2_. Following incubation, culture medium was aspirated and plates were gently washed by immersion in water to remove non-adherent cells. Plates were allowed to dry, after which 50 µL of 0.1% CV was added to each well and incubated for 15 min at room temperature. The CV solution was removed, plates were washed again to remove excess stain, and the plates were allowed to dry. Bound CV was solubilized by addition of 200 µL of 30% acetic acid. Extracted CV was diluted 1:4 in water in a new 96-well plate, and absorbance at 575 nm was measured using a Synergy H1 multimode plate reader with Gen5 microplate reader software (v 3.11). Unless otherwise indicated, biofilm assays were performed with four biological replicates, each measured in technical quadruplicate.

### Growth Measurements Under Acidic Starting Conditions

To assess strain-dependent growth under acidic conditions, overnight cultures were prepared and normalized as described above and inoculated into TYG. Immediately prior to inoculation, the starting pH of the medium was adjusted to pH 7.0, 6.0, or 5.5 using 1N HCl. Growth measurements were performed using a Bioscreen C MBR automated turbidometric analyzer. Optical density at 600 nm (OD_600_) was recorded every 0.5 h for 24 h, and wells were overlaid with 50 µL sterile mineral oil. Growth curves were used to calculate doubling time, final culture yield, and lag-associated growth. Final yield was determined from the mean of the final two OD_600_ readings, while lag was operationally defined as the time required for a culture to exceed OD_600_ = 0.1. Doubling times were determined from the exponential portion of the growth curve. Each assay was completed with three biological replicates with three technical replicates.

To evaluate retained biofilm biomass under the same acidic starting conditions, strains were inoculated into TYGS adjusted to pH 7.0, 6.0, or 5.5 and incubated for 24 h. Biofilm biomass was measured using the CV assay described above. Because isolates differed substantially in baseline biofilm formation, OD_575 nm_ values obtained at pH 6.0 and pH 5.5 were expressed as a percentage of the corresponding pH 7.0 value for each isolate. Isolates SMU041 and SMU053 were excluded from normalized comparisons because their low baseline biofilm biomass at pH 7.0 prevented meaningful normalization.

### Measurement of Acid-Associated Fluorescence Within Biofilms

To evaluate relative acid-associated fluorescence within mature biofilms, *S. mutans* isolates were grown as 18 h biofilms in TYGS as described above. Following incubation, biofilms were stained with pHrodo Green AM and Hoechst 33342 as a total-cell stain. pHrodo fluorescence increases under increasingly acidic conditions and was therefore used as a relative indicator of acid-associated microenvironments rather than as a calibrated measurement of absolute biofilm pH. Biofilms were imaged using a Lionheart FX automated microscope with a 40× objective, and z-stacks encompassing an approximately 256 × 256 µm field and 30-µm depth were collected. Relative pHrodo fluorescent intensity within individual microcolonies was measured using Gen5 Image+ software (v 3.11). Microcolonies from five independent image fields were analyzed for each isolate, and individual measurements were averaged to obtain isolate-level values for comparison with microcolony volume and other physiological phenotypes.

### Determination of Minimal Biofilm Inhibitory Concentration

The susceptibility of *S. mutans* isolates to chlorhexidine (CHX) and cetylpyridinium chloride (CPC) were assessed by measuring biofilm biomass formed during continuous exposure to each antimicrobial. Overnight cultures were normalized and inoculated into 96-well plates containing TYGS supplemented with increasing concentrations of CHX or CPC. The concentration series tested was 0.25-1.50 µg mL^-1^ and 0.25-4.00 µg mL^-1^, with untreated TYGS (0 µg mL^-1^) serving as the control. Biofilms were incubated for 24 h at 37°C in 5% CO_2_ and quantified using the CV assay described above. OD_575 nm_ values at each antimicrobial concentration were expressed as a percentage of the corresponding untreated control for each isolate. SMU041 and SMU053 were excluded from comparisons in which baseline CV accumulation was insufficient for meaningful normalization.

### Short-Term Chlorhexidine Exposure and Membrane Integrity

To evaluate the effects of a brief antiseptic challenge on established biofilms that would mimic usage of mouthwashes or rinses, seven *S. mutans* isolates representing distinct biofilm architecture and antimicrobial-response phenotypes were selected for analysis. Biofilms were grown for 18 h in TYGS and subsequently exposed for 0.5 minutes to either 50 or 500 µg mL^-1^ CHX. Treatment with 1× PBS served as the negative control, whereas 70% ethanol served as a positive membrane-compromise control. Following treatment, biofilms were immediately washed three times with PBS and stained with Hoechst 33342 and SYTOX Green (5 μM). Biofilms were imaged using a Nikon A1R confocal microscope. For single-cell analysis, NIS-Elements General Analysis (GA3), in combination with the existing NIS-Elements software AI tool, was utilized. Binary masks were generated from the Hoechst total-cell signal to define individual bacterial cell parameters, followed by quantification of SYTOX Green fluorescence within each segmented cell. A fixed fluorescent-intensity threshold was applied across all treatment conditions to classify cells as SYTOX positive. The percentage of SYTOX positive cells was calculated for each isolate and treatment condition.

### Statistical Analysis

Graphing and statistical analyses were performed using GraphPad Prism version 11.0. For comparisons of biofilm cellular biomass, glucan biomass, eDNA biomass, and microcolony volume between saliva-free and saliva-containing conditions, multiple unpaired *t* tests with Welch’s correction were performed. Differences in normalized CV biomass between acidic starting conditions were similarly evaluated using multiple unpaired *t* tests with Welch’s correction. Associations among isolate-average cellular biomass, glucan biomass, eDNA biomass, microcolony volume, and pHrodo fluorescent intensity were evaluated by simple linear regression, with coefficients of determination (R^2^) reported. For short-term CHX exposure experiments, differences in the percentage of SYTOX-positive cells among isolates and treatment conditions were analyzed by ordinary two-way analysis of variance (ANOVA) followed by Tukey’s multiple-comparisons test.

## Data Availability

The raw and processed data generated in this study have been deposited in FigShare and are publicly available under DOI **10.6084/m9.figshare.33289374**. Confocal microscopy image datasets generated in this study have been deposited in the BioImage Archive under accession number **S-BIAD4017**. Whole-genome sequence data for the isolate collection are available under the accession numbers previously reported in Cornejo et al (17).

## ACKNOLWEDGEMENTS

This work was supported by Colgate CARE Grant A-2023-5079-OC. Super-resolution confocal imaging was completed with resources from the Campus Microscopy and Imaging Facility (CMIF) and the OSU Comprehensive Cancer Center (OSUCCC) Microscopy Shared Resource (MSR), The Ohio State University. This facility is supported in part by grant P30 CA016058, National Cancer Institute, Bethesda, MD. We also acknowledge support from the Ohio State University Center of Microbiome Science and Infectious Diseases Institute through the Microbiome Platform within the Genomics and Microbiology Solutions Laboratory.

## AUTHOR CONTRIBUTIONS

Kyulim Lee: Investigation, Formal analysis, Writing – original draft.

Daniel I. Peters: Investigation, Formal analysis, Writing – review & editing.

Madisen Bangs: Investigation, Formal analysis, Writing – review & editing.

Delaney Hancock: Investigation, Formal analysis, Writing – review & editing.

Nicole Fleming: Investigation, Formal analysis, Writing – review & editing.

Jarett Pittman: Investigation, Formal analysis, Writing – review & editing.

Taylor Martinez: Investigation, Formal analysis, Writing – review & editing.

Alyssa N. Deever: Formal analysis, Data curation, Writing – review & editing.

Justin R. Kaspar: Funding acquisition, Project administration, Supervision, Writing – review & editing.

## ARTIFICIAL INTELLIGENCE (AI) UTILIZATION STATEMENT

The authors acknowledge the use of ChatGPT (GPT-5.6 Sol) in the final editing phases to assist with grammar polishing and improving the readability of the manuscript. The authors reviewed and edited the output and take full responsibility for the final content.

