## Supplementary material for "Strain-Level Diversity Decouples Biofilm Architecture, Acidogenic and Aciduric Traits, and Antimicrobial Tolerance in *Streptococcus mutans*": Figure S1

### SUPPLEMENTAL FIGURES AND FIGURE LEGENDS

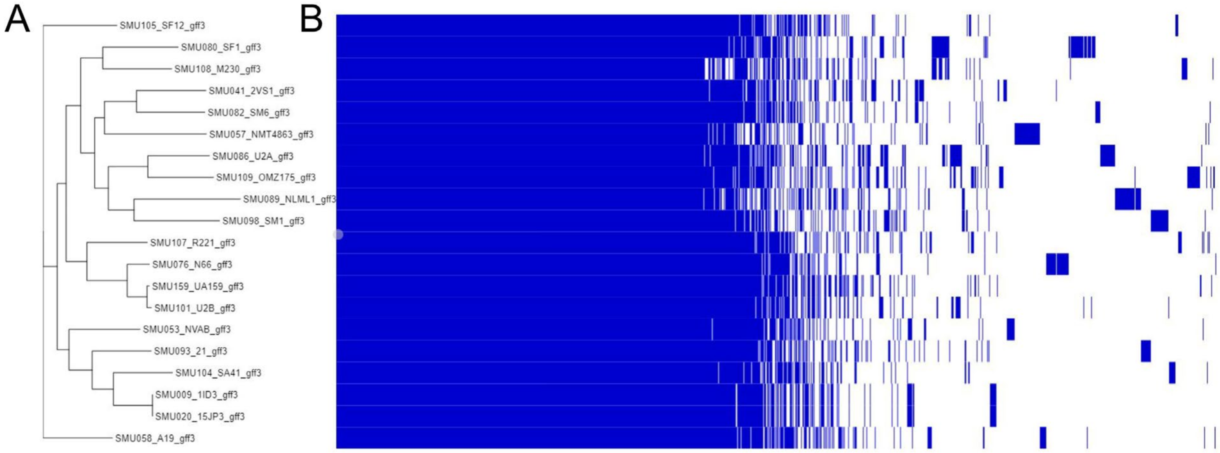


**Supplemental Figure 1. Genomic Diversity of *Streptococcus mutans* Clinical Isolates Used in this Project.** **(A)** Phylogenetic analysis of the 20 clinical isolate *S. mutans* strain panel used in this study using RAxML (Randomized Axelerated Maximum Likelihood). **(B)** Display of pangenome analysis using a Roary gene presence/absence matrix for 3353 total genes (gene pool of the 20 isolate strain panel). A blue line indicates gene presence that can be visually compared across the 20 strains. Isolates were chosen from a previously characterized panel (Palmer et al., 2013)and have been whole genome sequenced prior to this study (Cornejo et al., 2013).

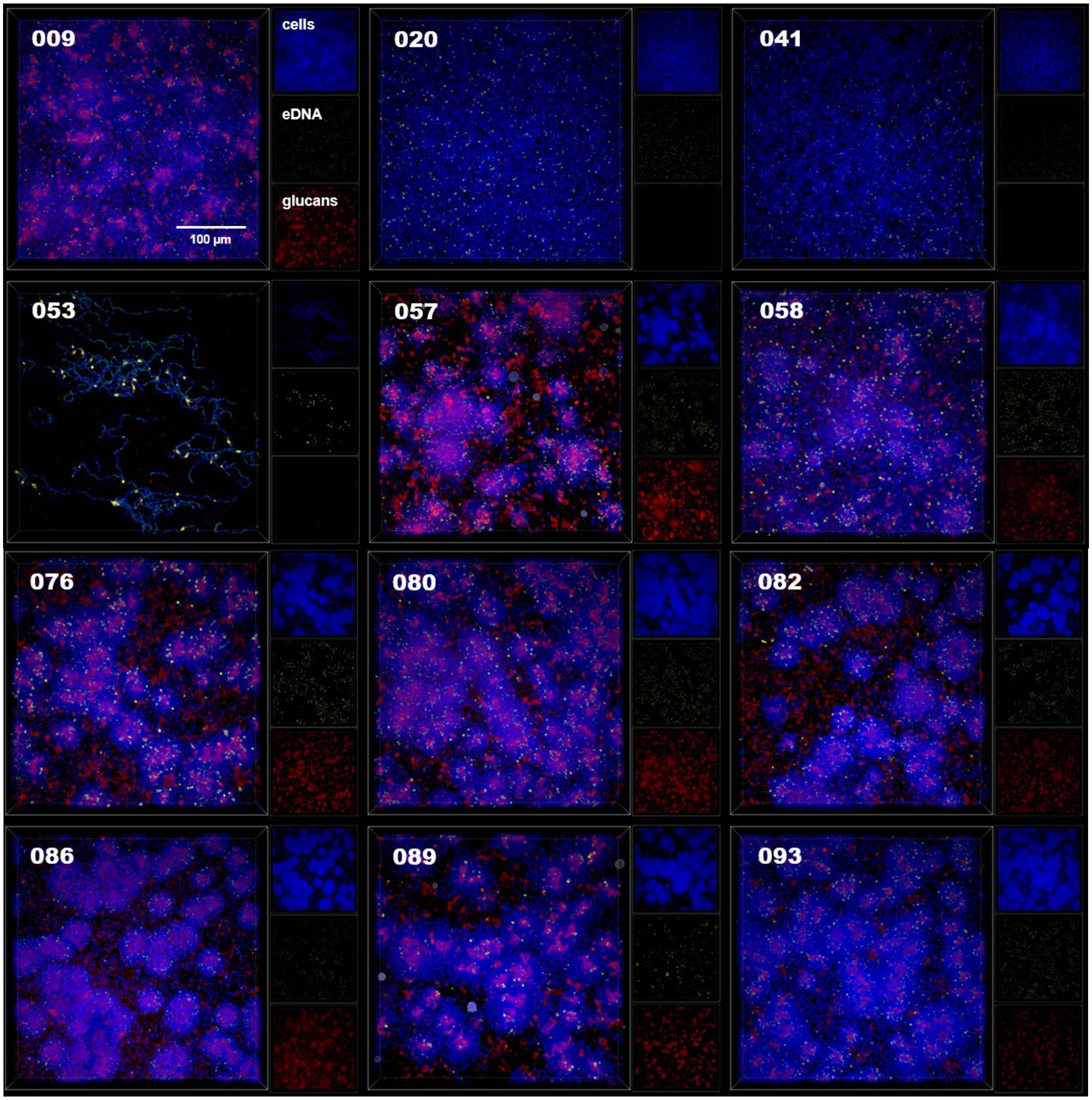
009
cells
eDNA
glucans
100 µm
020
041
053
057
058
076
080
082
086
089
093

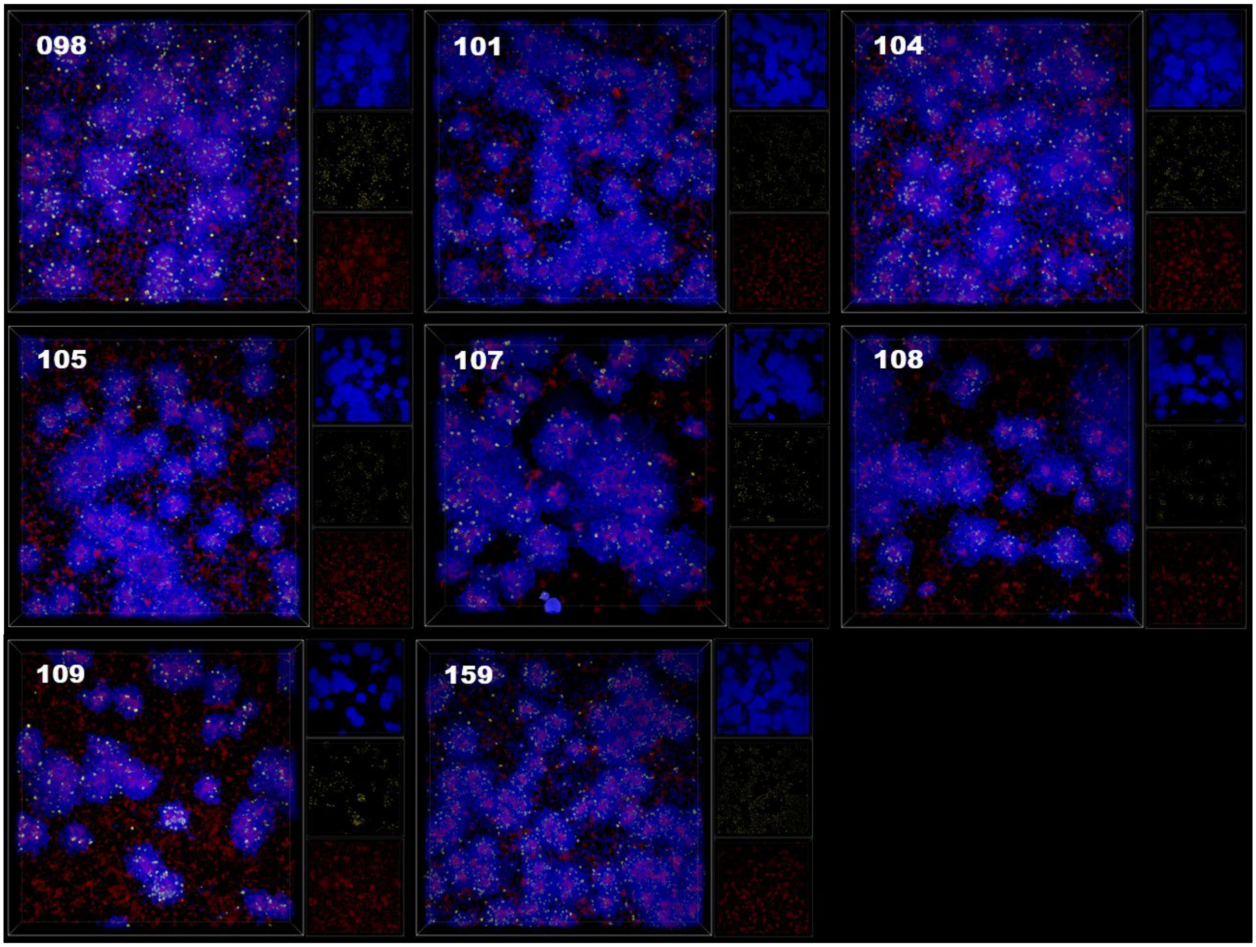


**Supplemental Figure 2. Microcolony Formation of 20 Diverse *Streptococcus mutans* Isolates.** Merged maximum intensity, 100x 3D models of a confocal-captured biofilm image oriented from the top down (Z+) of *S. mutans* monoculture biofilms, grown in tryptone-yeast extract medium with 10 mM glucose and 5 mM sucrose (TYGS). Biofilms were grown for 18 h. Individual channels recorded include DAPI (Hoechst total cell stain, [cells], blue), Texas Red (Alexa Fluor 594-labled α-dsDNA antibody, [eDNA], yellow), and CY5 (Alexa Fluor-647-labeled dextran [glucans], red) and are shown on the right side of the merged image. The strain identifier number (SMU###) is recorded in the top left of the merged image panel. Scale bar (100 µm) is shown in the bottom right merged image panel. Images are 256 µm (L) x 256 µm (W) x 30 µm (H).

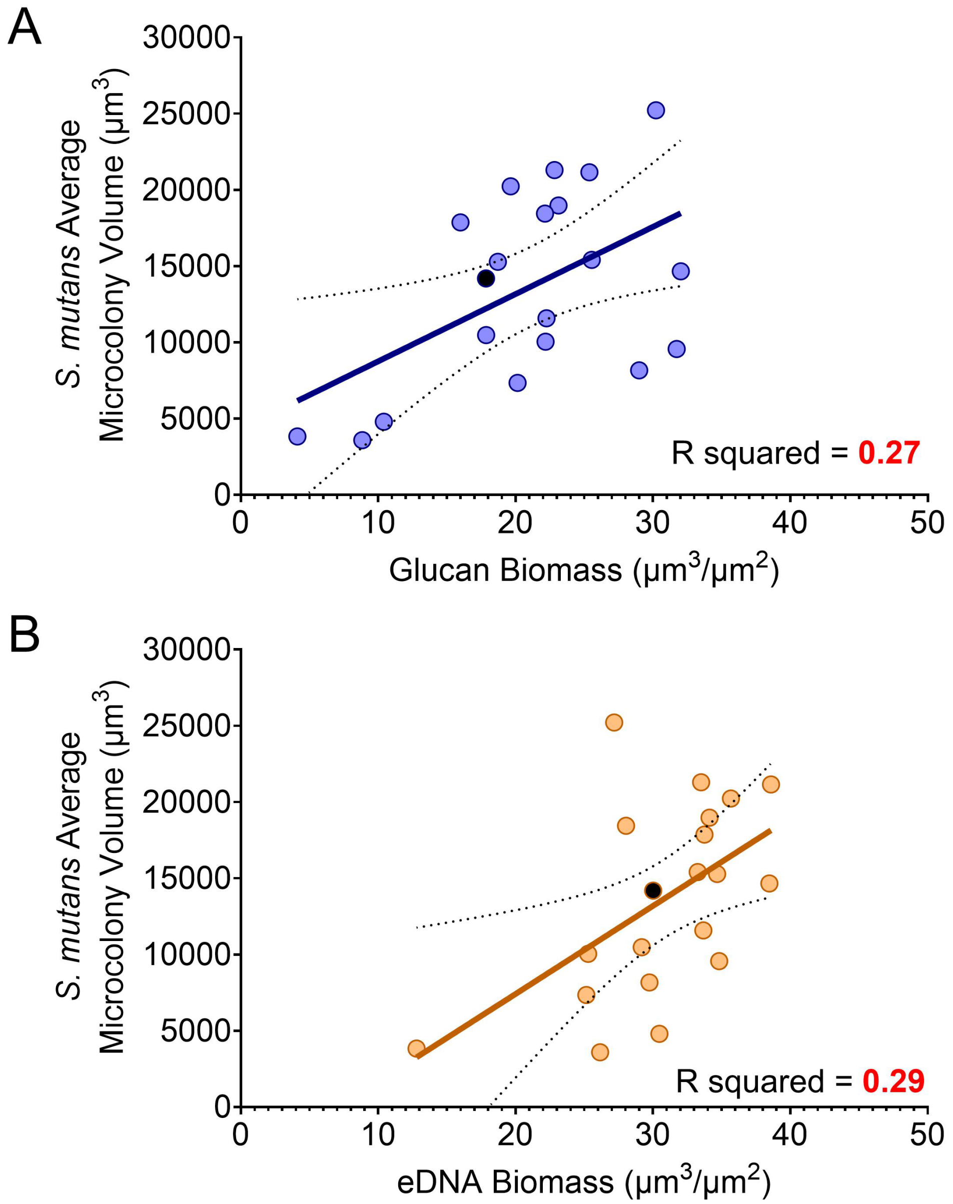


**Supplemental Figure 3. Association Between *Streptococcus mutans* Isolate Microcolony Volume and Matrix Production.** Simple linear regression plot of *S. mutans* isolate average microcolony volumes (y-axis) by either **(A)** average accumulated glucan biomass or **(B)** average accumulated eDNA biomass of that same isolate (x-axis). SMU159 (UA159) is represented by a black circle. Linear regression and R squared values were calculated using built-in analysis within GraphPad Prism (v11.0). Calculated R squared values are shown in bottom right corner.

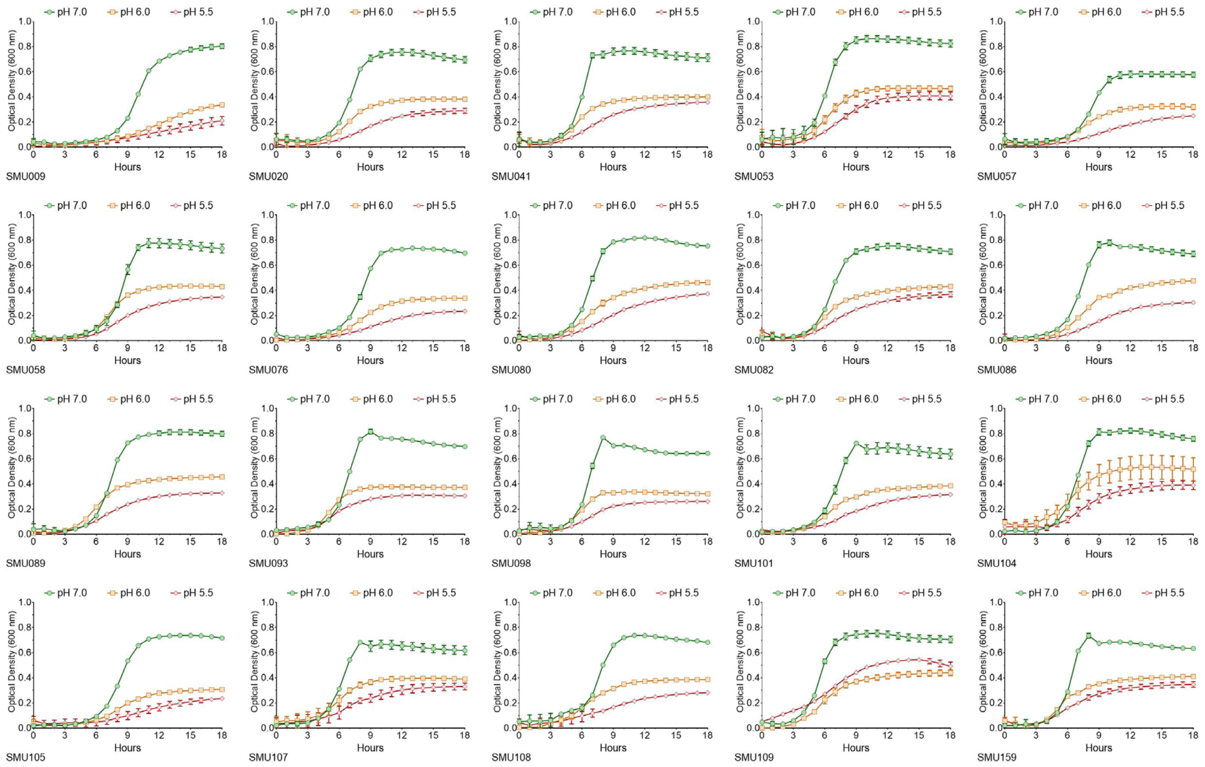


**Supplemental Figure 4. Growth Curves of Individual *Streptococcus mutans* Isolates with Varying Initial Medium pH Values.** Generated growth curves of individual *S. mutans* isolates (ID listed on bottom left) in TYG medium (10 mM glucose) with initial pH values of either 7.0 (green circles), 6.0 (orange squares), or 5.5 (red diamonds). Each strain was grown for 24 hours with readings recorded every 0.5 h (only every hour is plotted up to 18 h).

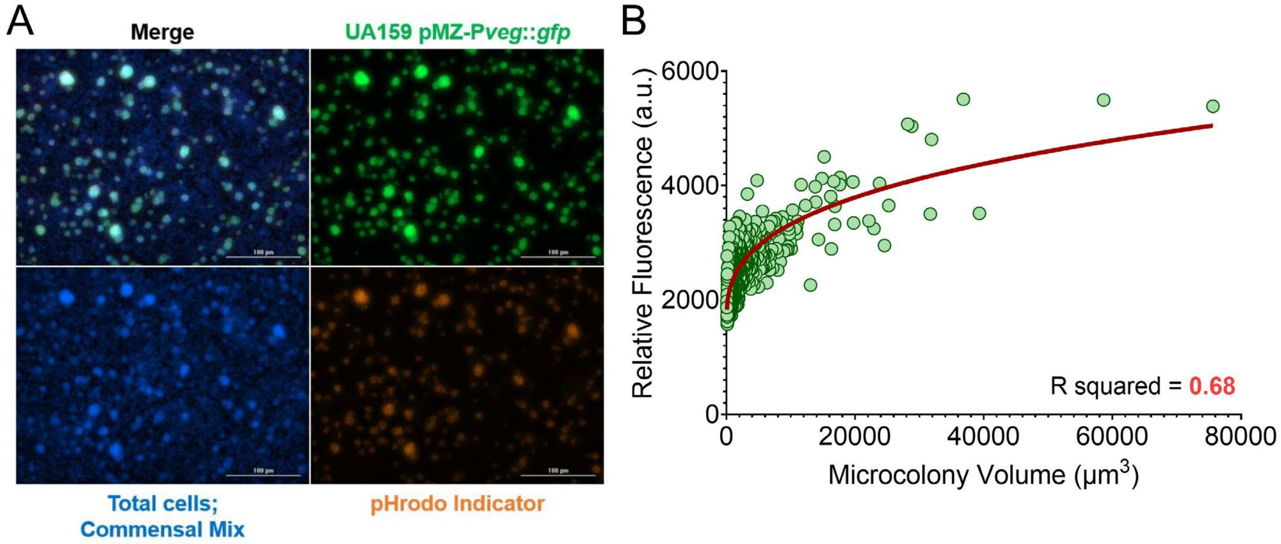


**Supplemental Figure 5. Specificity of pHrodo Dye to Acidic *Streptococcus mutans* Microcolonies.** **(A)** Maximum intensity Z-projection of 18 h *S. mutans* GFP-producing (UA159 pMZ-P*veg*::*gfp*) (Shields et al., 2019)cocultured biofilms with *Streptococcus gordonii* DL1. Merged channel is shown in the upper left, with individual channels recorded (additional panels) that include FITC (GFP – *S. mutans*, green, top right), DAPI (Hoechst total cell stain – *S. gordonii* and *S. mutans*, blue, bottom left) and RFP (pHrodo indicator, orange, bottom right). Scale bar is100 microns in the bottom right of the image. This experiment shows that the pHrodo indicator is specific to *S. mutans* microcolonies. **(B)** Plot of individual *S. mutans* microcolony measurements from the experiment, with individual microcolony volumes plotted on the x-axis and relative fluorescence of pHrodo indicator (shown as arbitrary units, a.u.) on the y-axis. Graphing and nonlinear regression completed in GraphPad prism software (v 11.0). n = 5 images quantified, 768 total *S. mutans* microcolonies analyzed.

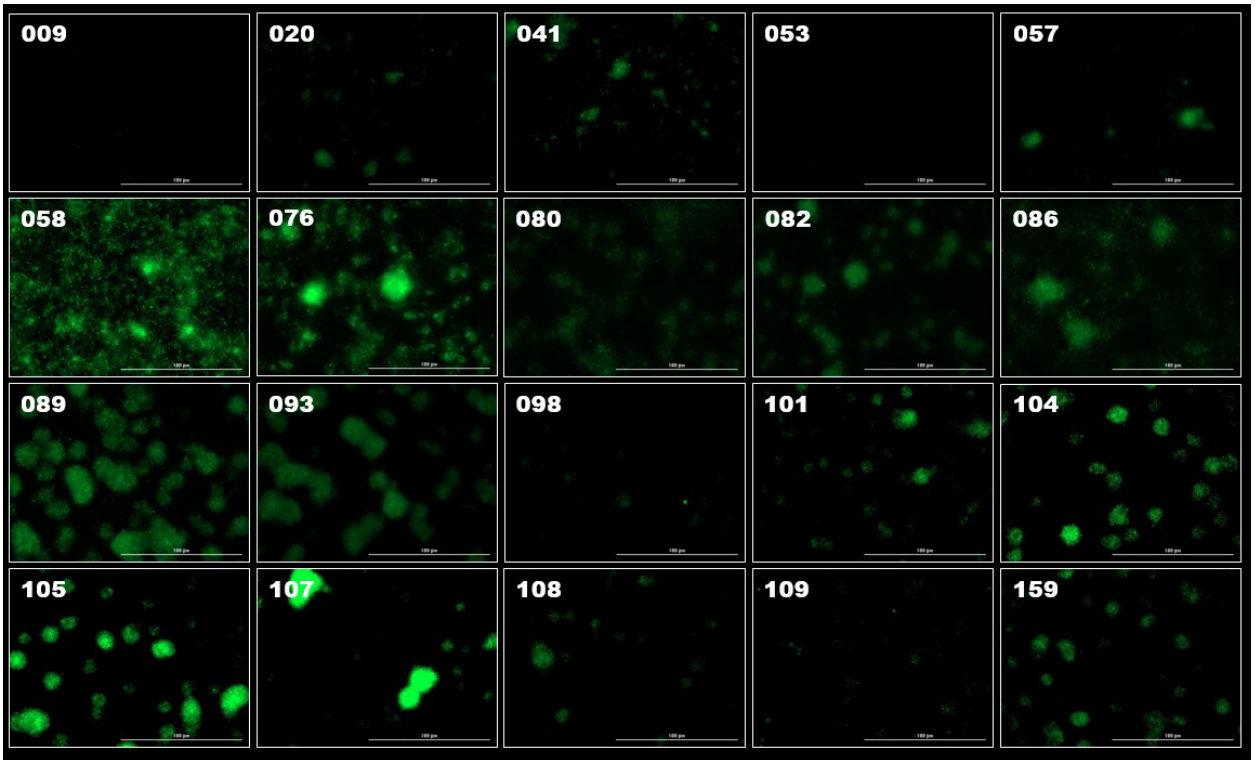


**Supplemental Figure 6. Detection of pHrodo within Microcolonies of Diverse *Streptococcus mutans* Isolates.** Maximum intensity 40x Z-projection images of 18 h *S. mutans* biofilms showing only the pHrodo Green AM pH indicator. These are the same images shown in Figure 5A (without the Hoechst total cell overlay). The isolate number (SMU###) for each representative image is shown in the top left. Scale bar (100 µm) is shown in the bottom right of the image. Images are 256 µm (L) x 256 µm (W) x 30 µm (H).

### SUPPLEMENTAL TABLES

**Supplemental Table 1. *Streptococcus mutans* isolate metadata and average microcolony volume ranking.**

| SMU Isolate #* | Original Strain Name* | Serotype* | Origin* | Average Microcolony Volume ($\mu m^3$) | Rank (by volume) |
|---|---|---|---|---|---|
| 009 | 1ID3 | e | Brazil | 8318 | 16 |
| 020 | 15JP3 | NA | Brazil | 3587 | 20 |
| 041 | 2VS1 | c | Brazil | 4698 | 18 |
| 053 | NVAB | c | USA | 3708 | 19 |
| 057 | NMT4863 | c | Japan | 20842 | 3 |
| 058 | A19 | c | UK | 10594 | 13 |
| 076 | N66 | c | UK | 21416 | 2 |
| 080 | SF1 | c | USA | 19070 | 6 |
| 082 | SM6 | c | Hong Kong | 7397 | 17 |
| 086 | U2A | e | Turkey | 16770 | 9 |
| 089 | NLML1 | e | UK | 9444 | 15 |
| 093 | 21 | e | Iceland | 20289 | 4 |
| 098 | SM1 | c | Hong Kong | 11679 | 12 |
| 101 | U2B | c | Turkey | 10518 | 14 |
| 104 | SA41 | c | South Africa | 17950 | 8 |
| 105 | SF12 | c | USA | 15236 | 10 |
| 107 | R221 | c | Brazil | 24803 | 1 |
| 108 | M230 | e | Brazil | 19188 | 5 |
| 109 | OMZ175 | f | NA | 18537 | 7 |
| 159 | UA159 | c | USA | 13723 | 11 |

* Metadata for the table was taken and previously published in Supplemental Material for (Cornejo et al., 2013)

### Supplemental Table 2. Sourcing of materials and supplies used in this study.

| Supplemental Table. Materials, Suppliers, Catalog Numbers, and Experimental Notes | | | |
|---|---|---|---|
| **Material** | **Supplier** | **Catalog Number** | **Notes** |
| **Bacterial Strains and Growth Conditions** | | | |
| Difco Brain Heart Infusion | Fisher Bioreagents | 237200 | Used for overnight growth; BHI also used for long-term storage medium containing 25% glycerol. |
| Difco Agar | Fisher Bioreagents | 214010 | Used with BHI to prepare agar plates for isolated colonies. |
| Glycerol | Fisher Bioreagents | BP229 | 25% final concentration in BHI for long-term storage at -80°C. |
| Tryptone | Fisher Bioreagents | BP1421 | 10 g/L in tryptone-yeast extract (TY) medium. |
| Yeast extract | Fisher Bioreagents | BP1422 | 5 g/L in TY medium. |
| Potassium phosphate dibasic ($K_2HPO_4$) | Sigma-Aldrich | P3786 | 3 g/L in TY medium. |
| D-(+)-Glucose | Sigma-Aldrich | G8270 | 10 mM in TYG; also present in TYGS biofilm medium. |
| Sucrose | Sigma-Aldrich | S7903 | 5 mM in TYGS for biofilm experiments. |
| **Human Saliva Preparation** | | | |
| Pooled human saliva | Innovative Research | IRHUSL250ML | Centrifuged at 4,500 rpm for 10 min; 0.22-µm filtered; stored as 10-mL aliquots; mixed 1:1 (v/v) with TY medium. |
| Millex 0.22 µm, diam. 33 mm, polyethersulfone (PES) membrane filter unit | Millipore | SLGPM33RS | Used to filter processed pooled human saliva. |
| **Biofilm Growth and Extracellular Matrix Labeling for Microscopy** | | | |
| Alexa Fluor 647-labeled dextran | Invitrogen | D22914 | 10,000 molecular weight (MW); anionic, fixable; 1 µM during biofilm growth to label glucan. |
| 12-well glass-bottom black plates | Cellvis | P12-1.5H-N | Used for biofilm growth and microscopy. |
| Bovine serum albumin (BSA) | Thermo Scientific | J61655.AK | 3% in PBS blocking buffer; 30 min at room temperature. |
| Anti-dsDNA murine monoclonal antibody (35I9 DNA) | Abcam | ab27156 | 2 µg/mL in BSA blocking buffer; 1 h at room temperature. |
| Alexa Fluor 594-labeled goat anti-mouse IgG secondary antibody | Invitrogen | A32742 | Highly cross-absorbed; 2 µg/mL; 1 h at room temperature. |
| Hoechst 33342 | Thermo Scientific | 62249 | 5 µM final concentration; 15 min; total-cell stain. |
| **Super-Resolution Confocal Microscopy and Image Analysis** | | | |
| A1R confocal microscope | Nikon | | 100X oil-immersion objective; 1.45 numerical aperture; 0.13-mm working distance; 30-µm z-depth; 0.5-µm steps. |
| NIS-Elements software (v6.02.03) | Nikon | | Biofilm image analysis software; Denoise.ai used uniformly before analysis. |
| BiofilmQ | https://drescherlab.org/data/biofilmQ/docs/ | | Microcolony segmentation; threshold >2,000 a.u.; minimum object size >5 µm. |
| **Crystal Violet Biofilm Biomass Quantification** | | | |
| 96-well, cell culture-treated, flat-bottom microplates | Fisherbrand | FB012931 | Used for crystal violet biofilm biomass retention assays. |
| Crystal violet | Fisher Chemical | C581 | 0.1%; 50 µL/well; 15 min at room temperature. |
| Acetic acid | RICCA Chemical | 1383032 | 30%; 200 µL/well to solubilize bound crystal violet. |
| Synergy H1 multimode plate reader | Agilent BioTek | | Absorbance measured at 575 nm. |
| Gen5 microplate reader software (v3.11) | Agilent BioTek | | Used with the Synergy H1 plate reader for absorbance measurements. |
| **Growth Measurements Under Acidic Starting Conditions** | | | |
| Hydrochloric acid (HCl) | Sigma-Aldrich | H1758 | Adjusted to 1 N; used to adjust starting medium TYG or TYGS pH to 7.0, 6.0, or 5.5. |
| Bioscreen C MBR automated turbidometric analyzer | Growth Curves Ab Ltd. | | OD600 nm recorded every 0.5 h for 24 h. |
| Mineral oil | Fisher Bioreagents | O121 | 50 µL/well overlay. |

| | | | |
|---|---|---|---|
| **Measurement of Acid-Associated Fluorescence Within Biofilms** | | | |
| pHrodo Green AM | Thermo Scientific | P35373 | Used as a relative indicator of acid-associated microenvironments in 18 hr biofilms. |
| Lionheart FX automated microscope | Agilent BioTek | | 40X objective; approximately 256 × 256 µm field; 30-µm z-depth. |
| Gen5 Image+ software (v3.11) | Agilent BioTek | | Used to quantify relative pHrodo fluorescence within individual microcolonies. |
| **Determination of Minimal Biofilm Inhibitory Concentration** | | | |
| Chlorhexidine (CHX) | Tokyo Chemical Industry (TCI) | C1254 | Continuous exposure during 24 hr biofilm growth; concentration series 0.25-1.50 µg/mL. |
| Cetylpyridinium chloride (CPC) | Tokyo Chemical Industry (TCI) | A5161 | Continuous exposure during 24 hr biofilm growth; concentration series 0.25-4.00 µg/mL. |
| **Short-Term Chlorhexidine Exposure and Membrane Integrity** | | | |
| Ethanol solution 70% v/v | Fisher Bioreagents | BP82011 | positive membrane-compromised (cell death) control |
| SYTOX Green | Thermo Scientific | S7020 | 5 µM; 15 min; membrane-integrity stain. |
| **Statistical Analysis** | | | |
| GraphPad Prism software (v11.0) | GraphPad Software | | Used for graphing, linear regression, t tests, two-way analysis of variance, and multiple-comparisons testing. |

### SUPPLEMENTAL METHODS

*Phylogenetic and Pangenome Analysis*.

Whole-genome sequence data (.fasta and .gff files) for the *S. mutans* isolates were obtained from (Cornejo et al., 2013) (Genbank accession numbers for each isolate are listed in Table S4 of their Supplemental Material). Exisiting annotations for each genome sequence were used, and phylogenetic relationships among isolates were determined using RAxML (v 8.2.13) (Stamatakis, 2014). Pangenome analysis was performed using roary (v 3.13.0) (Page et al., 2015) to identify core and accessory genome content across the isolate panel. All analysis was completed in Galaxy (Jalili et al., 2021). The resulting phylogenetic tree and distribution of core and accessory genome features are presented in Figure S1.

*Measurement of Acid-Associated Fluorescence Within Cocultured Biofilms.*

To validate localization of the pH-sensitive fluorescent signal within microcolonies, GFP-expressing *S. mutans* UA159 (Shields et al., 2019) was cocultured with *Streptococcus gordonii* DL1 at a 10:1 ratio (due to fast growth rate of *S. gordonii* DL1 compare to *S. mutans* UA159) in TYGS. Following 18 h of growth, biofilms were stained with pHrodo Red AM and Hoechst 33342 and imaged as described in the main Materials and Methods section. Fluorescence associated with individual *S. mutans* microcolonies was quantified and compared with microcolony volume using nonlinear regression (GraphPad Prism v11.0).
